# Expression and localization of NMDA receptor GluN2 subunits in dorsal horn pain circuits across sex, species, and late postnatal development

**DOI:** 10.1101/2025.10.22.683915

**Authors:** Katherine Griffiths, Jennifer Armstrong, Newton Martin, Clare Murray-Lawson, Estefania Oneil, Laurence S. David, Santina Temi, Jessica Parnell, Christopher Rudyk, Julia Bursey, Jeffrey L. Krajewski, Jeff S. McDermott, Annemarie Dedek, Ariel J. Levine, Baolin Li, Eve C. Tsai, Michael E. Hildebrand

## Abstract

Despite being essential mediators of pain processing, the molecular identity of N-methyl-D-aspartate receptor (NMDAR) subtypes in nociceptive dorsal horn circuits is poorly understood, especially between sexes and in humans. Given the importance of GluN2 subunits in shaping NMDAR function and plasticity, we investigated the expression and localization of specific GluN2 NMDAR variants in the dorsal horn of viable spinal cord tissue from male and female rodents and human organ donors. Analysis of single-cell/nuclei sequencing datasets and quantitative reverse transcriptase polymerase chain reactions (qRT-PCR) revealed that the GluN2A (*GRIN2A*) and GluN2B (*GRIN2B*) subunits are robustly expressed in dorsal horn neurons of mice, rats and humans, with moderate expression of GluN2D (*GRIN2D*). Immunohistochemistry (IHC) with antigen retrieval demonstrated that GluN2A, GluN2B, and GluN2D proteins are all preferentially localized to the superficial dorsal horn of both adult rats and humans, which is conserved between males and females. Surprisingly, we found that these GluN2 NMDAR subunits are enriched in the lateral superficial dorsal horn in rats but not in humans, while presynaptic and neuronal markers are symmetrically distributed across the rat mediolateral axis. A dramatic shift in localization of GluN2A to the lateral superficial dorsal horn was observed across later postnatal development (PD21-PD90) in both male and female rats, with a corresponding change in synaptic NMDAR currents. This discovery of changes in NMDAR subunit distribution during maturation and between species will shed light on the physiological roles of NMDARs and their potential as therapeutic targets for pain.

**SIGNIFICANCE STATEMENT:** We used complementary single-cell/nuclei analysis, immunostaining, quantitative reverse transcriptase polymerase chain reactions, RNAscope in situ hybridization, and electrophysiological approaches to compare the relative expression of N-methyl-D-aspartate receptor (NMDAR) GluN2 subunits in dorsal horn spinal cord pain circuits of mouse, rat, and human spinal cord tissue. Through these comparisons, we find that the transcripts and proteins of the GluN2A, GluN2B, and GluN2D NMDAR subunits are robustly expressed in superficial dorsal horn neurons, with conserved expression across sex but important differences in expression and localization patterns across late development and between species. These discoveries shed light on the physiological roles of NMDARs and their utility as potential therapeutic targets for pain.

## INTRODUCTION

The spinal cord is a critical site of somatosensory processing, and distinct cellular circuits within the superficial dorsal horn (SDH) are devoted to gating and transmitting nociceptive signals. In order to develop better treatments for pain, we need to further understand the underlying neurobiological mechanisms of spinal nociception in rodent preclinical models (Koch et al., 2018), but also extend this knowledge to human cells and circuits (Renthal et al., 2021). Recent translational efforts have uncovered that the expression and function of specific molecular determinants of pain processing can diverge dramatically between rodent and human peripheral sensory neurons (Sheahan et al., 2018; Papalampropoulou-Tsiridou et al., 2022; Zhang et al., 2024). Similar cross-species comparisons of the expression profiles of molecular mediators of dorsal horn excitability are therefore urgently needed to develop novel and effective spinal cord-targeting analgesic approaches.

Excitatory NMDA receptors (NMDARs) are key regulators of neuronal excitability and plasticity throughout the CNS, including within the SDH. We have recently identified differences in the function of synaptic NMDARs in rodent versus human lamina I SDH neurons (Dedek et al., 2024), but potential differences in the underpinning NMDAR subunit expression profiles within the SDH have yet to be explored. Moreover, whether these NMDAR subunit expression patterns change or remain constant across development remains poorly understood. Glutamate-activated NMDARs are tetrameric complexes containing two mandatory GluN1 subunits and different potential combinations of four genetically-encoded GluN2 subunits (GluN2A-D) (Paoletti et al., 2013). In juvenile male and female rats, the GluN2A, GluN2B and GluN2D NMDAR subunits are all found in the dorsal horn, with differences in the degree of localization to the SDH between subtypes (Temi et al., 2021). In terms of potential changes across early development, the canonical switch from GluN2B- to GluN2A-dominated synaptic NMDARs that occurs during the first three weeks of development in the rodent brain (Paoletti et al., 2013) is not observed in lamina II SDH neurons of male rats (Mahmoud et al., 2020). Studies that have investigated developmental changes in NMDAR subunit expression in SDH circuits have mainly been restricted to this early postnatal window (de Geus et al., 2020). Thus, whether NMDAR subunit expression and function changes across later postnatal development into adulthood needs to be tested, as this is critical for understanding changes in sensory processing, maladaptive plasticity (Brewer and Baccei, 2020), and the increased propensity for developing many forms of chronic pain across the lifespan (Domenichiello and Ramsden, 2019).

The dysregulation of SDH NMDARs that drives hyperexcitability and pathological pain can also vary by biological sex. We recently found that potentiation of GluN2B NMDARs by BDNF is restricted to male SDH neurons for both rodent and human pathological pain models (Dedek et al., 2022), a sexual dimorphism also found in activity-induced muscle pain (Hayashi et al., 2024). Given these findings, do NMDARs within the SDH diverge between sexes at baseline? Functionally, synaptic NMDAR responses in lamina I neurons are conserved between sexes in rats and humans (Dedek et al., 2024). However, the majority of studies on underlying NMDAR subunit expression have not been sex-inclusive, with only 1 of 13 studies identified from a recent systematic review on NMDAR expression including both sexes (de Geus et al., 2020). Considering sex as a biological variable will therefore be essential in investigating how spinal NMDAR subunit expression is conserved or diverges in basal physiological states.

Spinal NMDARs remain a promising molecular pain target (Hanson et al., 2024), and therefore need to be systematically characterized in the relevant SDH circuits across the key biological variables of development, sex and species. Here, we combine complementary approaches in mouse, rat, and human spinal cord tissues to address these translational gaps. We find that the transcripts and proteins of the GluN2A, GluN2B, and GluN2D NMDAR subunits are robustly expressed in SDH neurons, with conserved expression across sex but important differences in expression and localization patterns across late development and between rodents and humans.

## METHODS

### Ethical Approval

All rat experiments were approved by the Animal Care Committee at Carleton University, according to the guidelines and policies provided by the Canadian Council for Animal Care, Carleton University, the University of Ottawa Heart Institute. The collection and use of human tissue from organ donors were approved by the Ottawa Health Science Network Research Ethics Board (Protocol ID# 20150544-01H) and the Carleton University Research Ethics Board B (Ethics Protocol #104836). Informed written consent was obtained from the donor’s supplementary decision makers on behalf of the patients.

### Animals

Male and female Sprague-Dawley rats were used throughout this study at various developmental stages: juvenile (aged postnatal day 21; PD21), adolescent (aged postnatal day 42; PD42), and adult (aged postnatal day 90 and older; PD90+). All rats were provided by Charles River Laboratories. The rats were housed in same-sex pairs under a 12-hour light/dark cycle and had access to water and food *ad libitum*.

### Rat Spinal Cord Isolation and Preparation

Using an intraperitoneal injection of 3g/kg urethane (Sigma), the rats were deeply anesthetized before the spinal cord was isolated and removed via posterior laminectomy. Following dissection, the cords were promptly transferred into ice-cold oxygenated protective sucrose solution (50 mM sucrose, 92 mM NaCl, 17 mM D-glucose, 26mM NaHCO_3_, 5 mM KCl, 1.25 mM NaH_2_PO_4_, 0.5 mM CaCl_2_, 7 mM MgSO_4_, 1 mM kynurenic acid), bubbled with carbogen (95% O_2_, 5% CO_2_), as the meninges were removed, and the nerve roots were trimmed.

For IHC experiments, the tissue was immediately fixed in a 4% paraformaldehyde (PFA) solution in 0.1M phosphate buffer (PB) for 24 h at 4, then placed in a first wash of 10% sucrose solution in PB for 24 h, a second wash in 10% sucrose solution in PB for 6 to 24 h, and then a final wash in 30% sucrose solution in PB for at least 72 h, all performed at 4. Tissue sections were flash frozen in Cryomatrix (Fisher Scientific) using isopentane that was chilled with liquid nitrogen and stored at -80 until cryosectioning. The rat spinal cord tissue was sectioned in a transverse plane at 25μm using a microtome cryostat (Thermo Fisher Scientific) at -20 and was mounted immediately on pre-treated microscope slides (Fisherbrand^TM^ Superfrost^TM^ Plus) in a serial fashion and stored at -80 prior to IHC experiments.

For qRT-PCR experiments, spinal sections were mounted on a paraffin block and freeze sprayed (Fisherbrand^TM^ Super Friendly Freeze’It^TM^). A scalpel was used to cut the spinal cord in the horizontal plane at approximately the central canal to separate the tissue into dorsal and ventral horn segments. Dorsal horn segments were immediately placed on dry ice and transferred to a -80°C freezer for storage until use.

### Human Donors

Spinal cord tissue was collected from male (*n = 10*) and female (*n = 12*) adult human organ donors (aged 20-70, rounded to the nearest 5 for privacy reasons), the majority of whom were neurological determination of death (NDD) donors, with a smaller subset from donation after circulatory death (DCD), through the Trillium Gift of Life Network, as previously described (Dedek et al., 2019, 2022, 2025; Galuta et al., 2020; Parnell et al., 2023). All organ donor candidates were pre-screened to identify and exclude any communicable diseases (i.e., HIV/AIDS and syphilis) or neurological conditions (i.e., cancer, chronic pain, or spinal cord injuries) which would have interfered with the objectives of this study.

### Human Spinal Cord Isolation and Preparation

Prior to human tissue collection, hypothermia was induced using a cooling bed and neuroprotective magnesium-containing solution was typically perfused into the body. Transplant organs were harvested for donation, followed by a vertebral corpectomy to isolate and collect lumbar and/or thoracic regions of the spinal cord. This procedure was completed within 3 hours after aortic cross-clamping or organ flushing. The isolated tissue was immediately transferred into oxygenated, ice-cold sucrose cutting solution (composition detailed above), and meninges and blood vessels were quickly removed under a dissection microscope. Human donor tissue was processed using the same procedures described above for rat tissue for IHC (16 of 22 samples) and qRT-PCR (all samples). The remaining IHC samples were processed as free-floating tissue (see below).

### Free-Floating Human Tissue Preparation

A subset of human spinal cord tissue samples (*n = 6*) was processed as free-floating slices rather than being slide-mounted, as these samples were also served as controls for the experiments described in (Dedek et al., 2019). The isolation method used in the operating room followed the same protocol as outlined above; however, for these samples, the tissue was bubbled in saline for 70 minutes before being fixed in 4% PFA solution in PB for 24 to 36 h at 4□. The tissue was then placed in a first wash of 10% sucrose solution in PB for 24 h, a second wash in 10% sucrose solution in PB for 6 to 24 h, and then a final wash in 30% sucrose solution in PB for at least 72 h, all at 4□. Tissue samples were transferred directly into a cryoprotectant solution (28.7mL of 27.6g/L sodium phosphate monobasic [pH 7.3], 96.3mL of 28.4g/L sodium phosphate dibasic [pH 7.3], 375mL of 100μl/L diethylpyrocarbonate water, 300mL ethylene glycol, 200mL glycerol) at -20□ until sectioning. Using a Leica SM2000R microtome the tissue was sliced in transverse sections at 25μm and stored again in cryoprotectant solution at -20°C prior to slide mounting and IHC experiments. The free-floating (*n = 6*) and slide-mounted (*n =* 16) human tissue samples were combined into a single experimental group (*N =* 22) since the immunoreactivity patterns of GluN2A, GluN2B and GluN2D were conserved across both methodologies.

### Immunohistochemistry

IHC techniques were first used to investigate the relative expression and mediolateral localization of GluN2A, GluN2B, GluN2D protein subunits in the dorsal horn of lumbar and thoracic spinal cord of 10 male and 12 female adult human organ donors as well as the lumbar spinal cord (L4 and L5) in 6 male and 6 female adult rats, as previously performed for juvenile rats (Temi et al., 2021). Double labeling with calcitonin gene-related peptide (CGRP) was used, as CGRP is a well-established anatomical marker of the SDH, due to the high density of presynaptic CGRP-afferent fibers that selectively innervate rat and human laminae I and II (Tie-Jun et al., 2001; Eftekhari and Edvinsson, 2011; Shiers et al., 2021). All rat tissue was processed in triplicates or quadruplets for each animal, with male and female samples stained in tandem on the same day. Human tissue was not stained in triplicates; instead, a single sample per donor was included in the analysis and stained alongside rat tissue when possible.

After an initial set of 3x5 min washes in phosphate buffer saline (PBS), the tissue was then incubated in a peroxidase blocking solution (50% methanol, 48.2% PBS, and 1.8% hydrogen peroxide) for 30 min at room temperature, followed by another 3x5 min in PBS.

We then used a protease-induced antigen retrieval protocol on the tissue by treating the slides with pre-warmed pepsin at 37°C for 5 min. This pretreatment effectively unmasks the epitopes of interest primarily by breaking down protein cross-linkages that occur with aldehyde fixation, thereby enhancing the immunoreactivity of the GluN2 subunits within the postsynaptic density (Nagy et al., 2004). Immediately after, the sections were washed 3x5 min in PBS at room temperature and incubated at room temperature for 1 h in a blocking solution (5% normal goat serum (NGS), 0.3% Triton-X, 0.3% bovine serum albumin (BSA), 94.4% PBS). After the incubation period, the blocker on the tissue was then replaced with a solution containing primary antibodies diluted in the same blocking solution used previously (Table 1) against either GluN2A and CGRP, GluN2B and CGRP, GluN2D and CGRP, for approximately 24 h at room. These antibodies have been widely used to successfully label GluN2 protein subunits in numerous published experiments, including IHC, cytochemistry, and western blot analyses (Del Toro et al., 2010; Swanger et al., 2013; Atkin et al., 2015; Dzamba et al., 2015; Telezhkin et al., 2016; Bhandage et al., 2017; Temi et al., 2021). To assess cross-species validity of the antibodies, epitope conservation analyses were performed by aligning the immunogen sequences to the corresponding human protein sequences. These anaes demonstrated high conservation of the target regions. Specifically, the GluN2A immunogen sequence (aa 41–53) showed 92.3% identity with a single conservative substitution (Asn to Thr), GluN2B (aa 323–337) was fully conserved (100% identity), and GluN2D (aa 345–359) showed 92.9% identity with a single conservative substitution (Arg to Lys).

**Table 1.** List of antibodies and dilutions used for immunohistochemistry experiments.

| **Antibody** | **Dilution** | **Manufacturer** | **Catalog #** |
| --- | --- | --- | --- |
| Rabbit anti-GluN2A | 1:2000 | Alomone Labs | ACG-002 |
| Rabbit anti-GluN2B | 1:1000 | Alomone Labs | AGC-003 |
| Rabbit anti-GluN2D | 1:1000 | Alomone Labs | AGC-020 |
| Mouse anti-CGRP | 1:5000 | Sigma-Aldrich | C-7113 |
| Mouse anti-NeuN | 1:5000 | Abcam | ab104224 |
| Goat anti-mouse IgG (H + L) Alexa Fluor 647 | 1:1000 | Invitrogen | A-21235 |
| Goat anti-rabbit IgG (H + L) HRP | 1:500 | Invitrogen | 31460 |

The following day, tissue sections were washed 3x5 min in PBS and incubated in secondary antibodies (goat anti-mouse AlexaFluor 647 and rabbit anti-mouse immunoglobulin G conjugated with horseradish peroxidase (HRP) H + L; Table 1), diluted in the blocking solution for 2 h at room temperature. The remaining steps of the IHC protocol were performed in the absence of light to prevent the excitation of the fluorophores. Next, the tissue was washed 3x5 min in PBS and incubated in tyramide signal amplification (TSA) diluted 1:50 in the supplied amplification diluent (TSA® Plus Cyanine 3 System, AKOYA Biosciences, NEL744001KT). The TSA incubation period varied by tissue type and subunit: on rat tissue, it was 2.5 min for GluN2A, and 7.5 min for GluN2B and GluN2D, on human tissue, a 5 min incubation was used for all three subunits. The resulting enzymatic reaction facilitates the deposition of Cy3, enhancing fluorescent signal and improving the visualization of labeled protein complex (Brenner et al., 2004; Willcockson and Valtschanoff, 2008). Immediately after the incubation, the tissue was washed in PBS (3x5 min), then incubated for 5 min in a Hoechst 33258 solution in PBS (1:1000; Thermo Fisher #H3569) to stain all cell nuclei, followed by a final set of PBS washes (3x5 min). All slides were cover-slipped using Fluoromount^TM^ Aqueous Mounting Medium (Sigma-Aldrich, RI∼1.4) and Fisherbrand^TM^ Microscope #1.5 coverslips (#22-266882), sealed with clear nail polish within 24-48 h to prevent drying, and stored at -20 until imaging.

Negative control experiments were completed on both rat and human tissue by omitting the GluN2 primary antibodies (rabbit anti-GluN2A, -GluN2B or -GluN2D) on one slide per sample, while including only the mouse anti-CGRP primary antibody. Confocal images of controls revealed no evident background staining from secondary antibodies or tissue autofluorescence (Supplementary Fig. 1). To assess potential contributions of lipofuscin autofluorescence, a small subset of sections was treated with TrueBlack® Lipofuscin Autofluorescence Quencher; no change in signal-to-noise ratio was observed, suggesting minimal contribution of lipofuscin to the detected signal.

Two additional preliminary IHC studies were conducted using the same experimental protocol. First, to assess GluN2A expression relative to neuronal density, GluN2A and the neuronal marker NeuN (Mullen et al., 1992) (Table 1) were co-stained in thoracic spinal cord tissue from adult rats (males: *n = 6*; females: *n = 6*). Second, to examine GluN2A localization across development, GluN2A and CGRP were co-stained in tissue from juvenile rats (PD21) and late adolescent (PD42) rats (*n = 4* per sex per age group), processed in parallel.

### Image Acquisition

All confocal z-stack compressed tiled images were acquired using a Zeiss Airyscan 800TM confocal microscope using a 20x objective lens and further processed using ZEN 3.1 imaging software. Identical microscope settings were applied to all images to ensure consistency, and laser intensity was appropriately selected to minimize photobleaching of the sample. To account for multiple tissue layers, stacks of horizontal plane images and orthogonal projections were acquired for each image of the right and left dorsal horns.

### Image Analysis

For rats, the L4 and L5 spinal cord segments were used for analysis. As previously described (Temi et al., 2021), prior to imaging, these cross sections were identified under a white light microscope, using a rodent spinal cord atlas to reference key cytoarchitectural features (Sengul et al., 2012). These included the anatomical characteristics of gray matter and dorsal column white matter, as well as the outer diameter of the spinal cord specific to L4 and L5 sections. Human analyses included both the thoracic and lumbar spinal segments due to variability in tissue availability across donors. The specific segment obtained depended on what was accessible at the time of organ retrieval and was initially identified by manual counting of vertebral levels. Given the inherent biological variability associated with human post-mortem tissue, segmental identity was subsequently confirmed using a standardized human spinal cord atlas (Sengul et al., 2012). For all experiments, we selected either the right side or the left side of the dorsal horn for analysis, prioritizing the side free from folds, damage or artifacts. In cases where both sides were viable, the images were examined for any instances of poor tiling during confocal image acquisition. If both the left and right dorsal horns were suitable for analysis, the representative side was selected at random using a coin flip (heads = right dorsal horn, tails = left dorsal horn).

Using FIJI ImageJ software, the SDH was defined by manually outlining the dorsal horn area of maximal CGRP-positive staining in both rat and human spinal sections. This tracing was performed exclusively on the CGRP-stained image, without viewing the Cy3-stained image (representing GluN2 subunits), to eliminate any potential experimenter bias in delineating the SDH region. A second trace was made to outline the DDH by referencing anatomical markers in the spinal cord tissue as indicated by a spinal cord atlas (Sengul et al., 2012). In rats, this DDH boundary extended to the central canal, whereas in humans it only extended approximately halfway down towards the central canal due to image acquisition constraints. The exact tracings of the SDH and DDH were then overlaid onto the Cy3-stained GluN2 subunit images. The pixel intensity per area was quantified for each outlined region, representing the expression of GluN2A, GluN2B, and GluN2D subunits. All values were then normalized to the pixel intensity of a small rectangular selection from the background (BG) in the white matter, medial to the dorsal horn, where Cy3 fluorescence was minimal Cy3. In our secondary study investigating NeuN protein expression, we did not have CGRP staining to act as a definitive marker for the SDH. Instead, the tracing was conducted using a rat spinal cord atlas as a reference (Sengul et al., 2012) with a clean curved line outlining the most ventral border of the SDH.

To quantify the expression of GluN2 subunits across the mediolateral axis of the SDH, the same CGRP-positive staining outlining the SDH used in the previous analysis was highlighted again. A 60 μm thick line was manually drawn following the natural curve of the SDH starting from the most medial region to the most lateral region, varying in length depending on the species (rats = 900 μm; humans = 1200 μm; Fig. 8A). ImageJ software quantified the pixel intensity of GluN2A, GluN2B and GluN2D subunit expression at 1.24 μm increments across the line. To reduce variability in staining intensity across images, the raw data was normalized to the average pixel intensity from a medial 0 μm to 50 μm range representing the lowest pixel intensity across the entire line scan. The mean pixel intensity across the medial SDH (150 μm to 350 μm in rats; 200 μm to 400 μm in humans) and the lateral SDH (600 μm to 800 μm in rats; 1000 μm to 1200 μm in humans) was calculated from each sample’s representative line scan. For all immunohistochemical analyses, normalized pixel intensity values in rats were averaged across three technical replicates (slices) per animal, whereas a single representative slice was analyzed per human donor. Statistical comparisons were performed at the level of the biological replicate, with animals or human donors treated as the independent experimental units. Sample sizes were as follows: rats, *n* = 6 animals per sex for each GluN2 subunit; humans, *n* = 12 female and 10 male donors per GluN2 subunit.

### Mouse and Human Single-Cell RNA Sequencing Data Information and Analysis

Processed single-cell and nuclei RNA sequencing data from both mouse and human samples were generously provided by Dr. Ariel Levine and Dr. Kaya Matson at the National Institute of Neurological Disorders and Stroke. The mouse data were sourced from a meta-analysis of six single-cell/nuclei RNA sequencing studies on lumbar spinal cord tissue (Russ et al., 2021). The human spinal cord data were derived from highly viable lumbar spinal cord tissue from four male and three female donors, aged approximately 20 to 80 years (Yadav et al., 2023).

We used a modular low-code pipeline, GScpter (https://github.com/NewtontheNeuron/GScpter) and the methods previously described in (Parnell et al., 2023) to analyze the gene expression of *GRIN1* (GluN1), *GRIN2A* (GluN2A), *GRIN2B* (GluN2B), *GRIN2C* (GluN2C), and *GRIN2D* (GluN2D) in defined dorsal horn neuronal clusters from both mouse and human single-cell RNA sequencing data (Russ et al., 2021; Yadav et al., 2023). Dot plots displaying the average expression and percent expressed of *GRIN1* and *GRIN2A-D* in the mouse and human dorsal horn neuronal clusters were generated using GScpter (https://github.com/NewtontheNeuron/GScpter).

### Quantitative Reverse Transcriptase Polymerase Chain Reactions

RNA extraction and complementary DNA (cDNA) synthesis were performed using the previously described procedure (Bellavance et al., 2024), with minor modification. Briefly, total RNA was extracted from rat and human dorsal horn spinal cord tissue using the RNeasy® Plus Mini Kit with QIAshredder columns (QIAGEN). Human tissue was pulverized in liquid nitrogen using a mortar and pestle with mechanical grinding into powder. Approximately 30mg of powdered tissue was homogenized in 600 uL Buffer RTL Plus. Rat tissue was homogenized directly in 600 uL of Buffer RTL Plus using mechanical disruption with a needle and syringe. Lysates were passed through QIAshredder columns (14,500 rpm, 3 min), followed by genomic DNA removal using gDNA Eliminator spin columns (11,000 rpm, 30 s). After addition of 70% ethanol, samples were applied to RNeasy spin columns and processed according to the manufacturer’s wash protocol (RW1 and RPE buffers). RNA was eluted in 50 µL RNase-free water and stored at −80 °C until use. cDNA was synthesized from 50 ng total RNA using the iScript^TM^ cDNA Synthesis Kit (Bio-Rad) in a 50 μL reaction volume. Reverse transcription was performed on a thermocycler under the following conditions: 25°C for 5 min, 46°C for 20 min, 95°C for 1 min, and hold at 4°C.

384-well qRT-PCR was performed using the SsoADVANCED^TM^ Universal SYBR® Green Supermix (Bio-Rad) on a CFX Real-Time PCR Detection System (Bio-Rad). Reactions (20 µL total volume) consisted of 9 µL master mix and 1 µL cDNA with gene-specific primers. All reactions were run in duplicate with no-template and no-RT controls. Cycling conditions were: 95 °C for 2 min (activation), followed by 40 cycles of 95 °C for 5 s and 60 °C for 30 s. Melt curve analysis (65–95 °C, 0.5 °C increments) was performed to confirm amplification specificity. β-Actin (rat: rActb; human: rACTB) served as the housekeeping gene. Rat primers included *Grin1*, *Grin2a*, *Grin2b*, *Grin2c*, *Grin2d*, *Grin3a*, and *Grin3b* (Table 2). Human primers included *GRIN1*, *GRIN2A*, *GRIN2B*, *GRIN2C*, *GRIN2D*, *GRIN3A*, and *GRIN3B* (Table 3).

**Table 2.** List of *Grin* primers (Bio-Rad) used for rat qRT-PCR experiments.

| Gene Target | Unique Assay ID | Assay Design |
| --- | --- | --- |
| *Grin1* | qRnoCID0007266 | Intron-spanning |
| *Grin2a* | qRnoCID0003413 | Intron-spanning |
| *Grin2b* | qRnoCID0002142 | Intron-spanning |
| *Grin2c* | qRnoCID0002895 | Intron-spanning |
| *Grin2d* | qRnoCID0009406 | Intron-spanning |
| *Grin3a* | qRnoCID0003085 | Intron-spanning |
| *Grin3b* | qRnoCED0004604 | Exonic |

**Table 3.** List of *GRIN* primers (Bio-Rad) used for human qRT-PCR experiments.

| Gene Target | Unique Assay ID | Assay Design |
| --- | --- | --- |
| *GRIN1* | qHsaCED0047208 | Exonic |
| *GRIN2A* | qHsaCID0007534 | Intron-spanning |
| *GRIN2B* | qHsaCID0015234 | Intron-spanning |
| *GRIN2C* | qHsaCID0016723 | Intron-spanning |
| *GRIN2D* | qHsaCID0016959 | Intron-spanning |
| *GRIN3A* | qHsaCED0048368 | Exonic |
| *GRIN3B* | qHsaCED0003359 | Exonic |

Rat (*n* = 16 total; 4 female lumbar, 4 male lumbar, 4 female thoracic, 4 male thoracic) and human (*n* = 16 total; 3 female lumbar, 3 male lumbar, 5 female thoracic, 5 male thoracic) experiments were run on single 384-well plates to enable direct quantitative comparisons. Melt curves were inspected to confirm single-product amplification. Cycle threshold (Ct) values were first averaged across technical duplicates and normalized to the plate average of the housekeeping gene *ACTB* using the following equation:

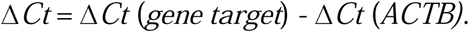

Relative expression was calculated and plotted as comparative Ct values using the following equation:

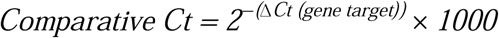

All analyses for RT-qPCR took place in Microsoft Excel and RStudio, using base, tidyverse, stats, gridExtra, purrr, dplyr, and ggplot2 packages. Statistical analyses were performed on ΔCt values to preserve the log-transformed scale.

### RNAscope Assay

Lumbar spinal cord tissue was collected from female rats (3 weeks and 14 weeks old) and embedded in O.C.T. compound. Tissues were sectioned 10 μm thick and mounted on Fisherbrand^TM^ Superfrost^TM^ Plus Gold microscope slides. To increase adherence, slides were air dried at -20°C in the cryostat for 1 h prior to storage, then stored at -80°C overnight or until ready to use. Frozen tissue slides were removed from -80°C and immediately fixed in 4% PFA in 1X PBS at room temperature for 30 min, and then rinsed in PBS for 1 min. To dehydrate the tissue, the slides were place successively in 50%, 70%, 100%, 100% ethanol baths at room temperature for 5 min each. RNAscope^TM^ assay (Wang et al., 2012) was performed using a commercially available kit and RNAscope^TM^ LS 2.5 probes against rat GluN2A subunits (*Grin2a*, NM_012573.3, bp2812–4265, ACD# 414628) from Advanced Cell Diagnostics (ACD). RNAscope hybridization was carried out using a RNAscope^TM^ LS 2.5 Multiplex Fluorescent assay (ACD# 322800) in combination with a Leica Biosystems BOND RX Automated Stainer to process samples according to the manufacturer’s (ACD) instructions. Positive controls were evaluated using RNAscope^TM^ LS 2.5 Multiplex Human positive control probe for fluorescent assay- against POLR2A (C1 channel) and PPIB (C2 channel), UBC (C3 channel). Negative control to establish background was evaluated by using RNAscope ^TM^ 3-plex LS Multiplex Negative control probe to DapB (of Bacillus subtilis strain). Dehydrated slides were placed in the Stainer to be pre-treated, including a protease IV treatment followed by the addition and hybridization of the desired probes with three signal amplifications. Fluorophores using Opal dyes were applied to visualize signal from each probe using the Vectra Polaris^TM^ Automated Quantitative Pathology Imaging System (AKOYA Biosciences).

### Electrophysiology Recordings on Lamina II Spinal Cord Neurons

Parasagittal spinal cord slices from male juvenile rats aged PD20-PD22 (*n =* 5 neurons) and adult rats aged PD90+ (*n =* 11 neurons) were visualized under a Zeiss microscope, as previously reported (Hildebrand et al., 2014; Mahmoud et al., 2020). Outer lamina II neurons were identified based on their location in the substantia gelatinosa. The external recording solution consisted of an artificial cerebral spinal fluid (aCSF) containing: 125 mM NaCl, 3 mM KCl, 26 mM NaCO_3_, 1.25 mM NaH_2_PO_4_, 2 mM CaCl_2_, 1 mM MgCl_2_, and 20 mM D-(+)- glucose, 10 μM bicuculline (Tocris Bioscience), 10 μM strychnine, 10 μM Cd^2+^, and 0.5 μM TTX (Alomone Labs). Patch-clamp recording borosilicate glass pipettes with typically resistances of 5 to 11 M Ω. were pulled using a Sutter Flaming micropipette puller and fire-polished. Pipettes were filled with an internal solution containing: 105 mM D-gluconic acid, 105 mM CsOH, 17.5 mM CsCl, 10 mM HEPES, 10 mM BAPTA, 2 mM Mg-ATP, 0.5 mM Na_2_-GTP (pH = 7.25; 295 mOsm). Using a whole-cell patch-clamp configuration, voltage-clamp recordings were obtained from neurons at room temperature using a MultiClamp 700B amplifier (Molecular Devices, Sunnyvale, CA, USA), Digidata® 1550 Data Acquisition System (Molecular Devices) and a personal computer running pClamp® 10.7 software. Initial recordings were obtained at −60 mV. Following 120 s of recording at −60 mV, the voltage was gradually increased to +60 mV to relieve the magnesium blockade of NMDAR responses. A minimum of 20 mEPSC events were recording at + 60 mV during a 10-min baseline recording period for each recorded lamina II neuron. All detected individual mEPSCs were manually aligned from their rising slope and averaged, as previously described (Hildebrand et al., 2014; Dedek and Hildebrand, 2024).

### Statistical Analysis

All statistical tests were completed using IBM® SPSS® Statistics software or RStudio® using base, tidyverse, stats, gridExtra, purrr, dplyr, and ggplot2 packages. Data are presented as mean ± standard error of the mean (SEM). Assumptions of normality and homogeneity of variance were assessed using the Shapiro–Wilk test and Levene’s test, respectivel, and were further evaluated by inspection of normality plots. When parametric assumptions were met, paired or independent-samples t-tests and two-way analyses of variance (ANOVAs) were used to assess group differences (see Supplementary Table 1 for summary of all statistical tests and results). Bonferroni-corrected post-hoc tests with bootstrapping were applied where appropriate. For all analyses, *p* <0.05 was considered statistically significant.

## RESULTS

### Immunostaining of GluN2A, GluN2B, and GluN2D NMDAR Subunits in the Dorsal Horn of Male and Female Adult Rats and Humans

To investigate the spatial expression of GluN2 NMDAR subunits across the dorsal horn in adult rats (PD90+), we co-stained lumbar spinal sections with selective antibodies against specific GluN2 subtypes as well as a marker of the SDH – calcitonin gene-related peptide (CGRP) (Parnell et al., 2023). Qualitative analysis revealed robust immunoreactivities for all three GluN2 subunits (GluN2A, GluN2B, and GluN2D) within the spinal dorsal horn, with an enhanced localization to the SDH and similar patterns of distribution for each subunit between female (Fig. 1-3A) and male (Fig. 1-3B) adult rats. In both sexes GluN2A labeling was localized to a combination of dense synaptic neuropil and somatic labeling throughout the dorsal horn (Fig. 1<u>(iv)</u>). The diverse subcellular distribution of GluN2B immunoreactivity closely mirrored that of GluN2A for both females and males (Fig. 2<u>(iv)</u>). Lastly, GluN2D subunit immunoreactivity appeared to be more moderate compared to GluN2A and GluN2B, with a potentially greater localization to the synaptic neuropil rather than cell bodies (Fig. 3<u>(iv)</u>). Positive staining for GluN2A, GluN2B and GluN2D was also detected in the dorsal horn white matter (Fig.1-3(iii)), suggesting that these NMDAR isoforms are expressed in cells and/or myelin beyond the dorsal horn gray matter. This localization indicates their presence in glial cells, ascending sensory pathways, and/or descending modulatory tracts, highlighting broader potential roles for these NMDAR subunits in spinal cord signaling and modulation (Bourinet et al., 2014).

**Figure 1.**
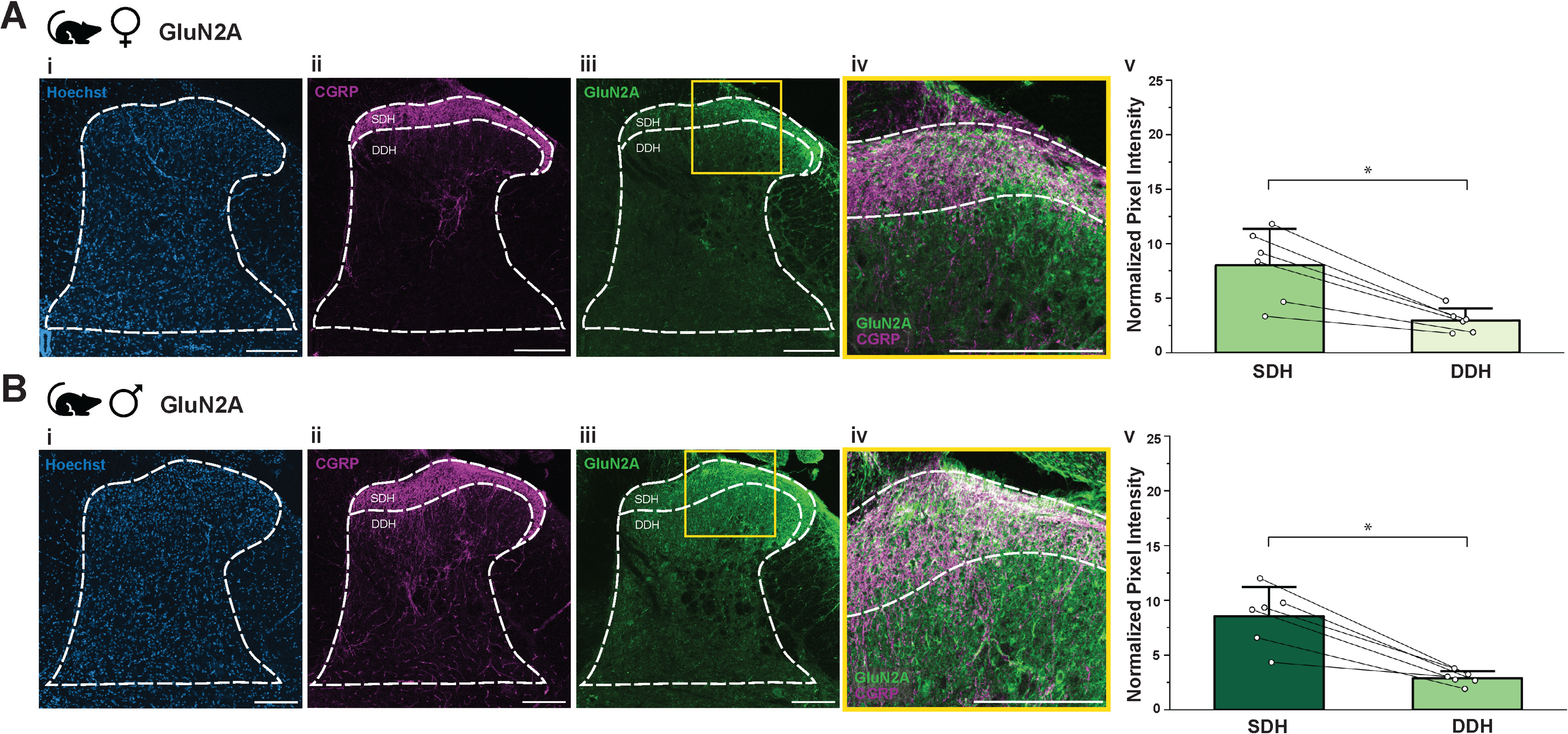
GluN2A subunits are significantly localized to the SDH compared to the DDH of the spinal dorsal horn in male and female adult rats. Representative confocal images (20× objective) from **(A)** female (L4) and **(B)** male (L4) adult rat dorsal horns showing (i) Hoechst nuclear staining to enable outlining of the dorsal horn grey matter (white dashed line) (ii) CGRP immunoreactivity marking the superficial dorsal horn (SDH; labelled, with additional marked line) as well as the remaining deeper dorsal horn (DDH), and (iii) GluN2A immunolabeling, with a yellow box highlighting the inset region shown in the panel to the right. (iv) Higher magnification images highlight GluN2A and CGRP colocalization within the SDH. Scale bars = 200 µm. (v) Quantitative analysis comparing the average expression (normalized pixel intensity) of GluN2A subunit immunoreactivity in the SDH versus the DDH of female (*n = 6* animals) and male (*n = 6* animals) adult rats aged postnatal day 90+. Data represents means ± SEM. *p<0.05.

**Figure 2.**
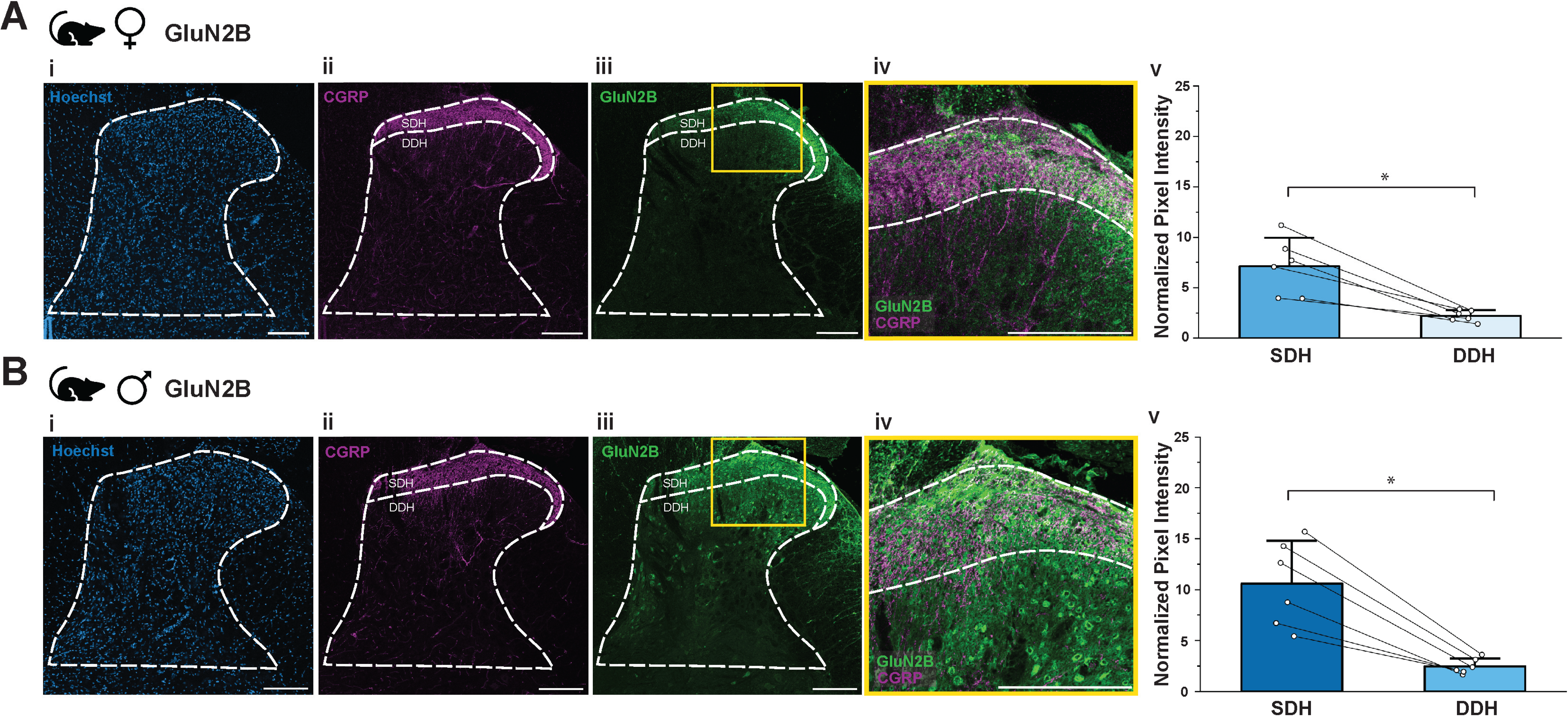
GluN2B subunits are significantly localized to the SDH compared to the DDH of the spinal dorsal horn in male and female adult rats. Representative confocal images (20× objective) from **(A)** female (L5) and **(B)** male (L4) adult rat dorsal horns showing (i) Hoechst nuclear staining to enable outlining of the dorsal horn grey matter (white dashed line) (ii) CGRP immunoreactivity marking the superficial dorsal horn (SDH; labelled, with additional marked line) as well as the remaining deeper dorsal horn (DDH), and (iii) GluN2B immunolabeling, with a yellow box highlighting the inset region shown in the panel to the right. (iv) Higher magnification images highlight GluN2B and CGRP colocalization within the SDH. Scale bars = 200 µm. (v) Quantitative analysis comparing the average expression (normalized pixel intensity) of GluN2B subunit immunoreactivity in the SDH versus the DDH of female (*n = 6* animals) and male (*n = 6* animals) adult rats aged postnatal day 90+. Data represents means ± SEM. *p<0.05.

**Figure 3.**
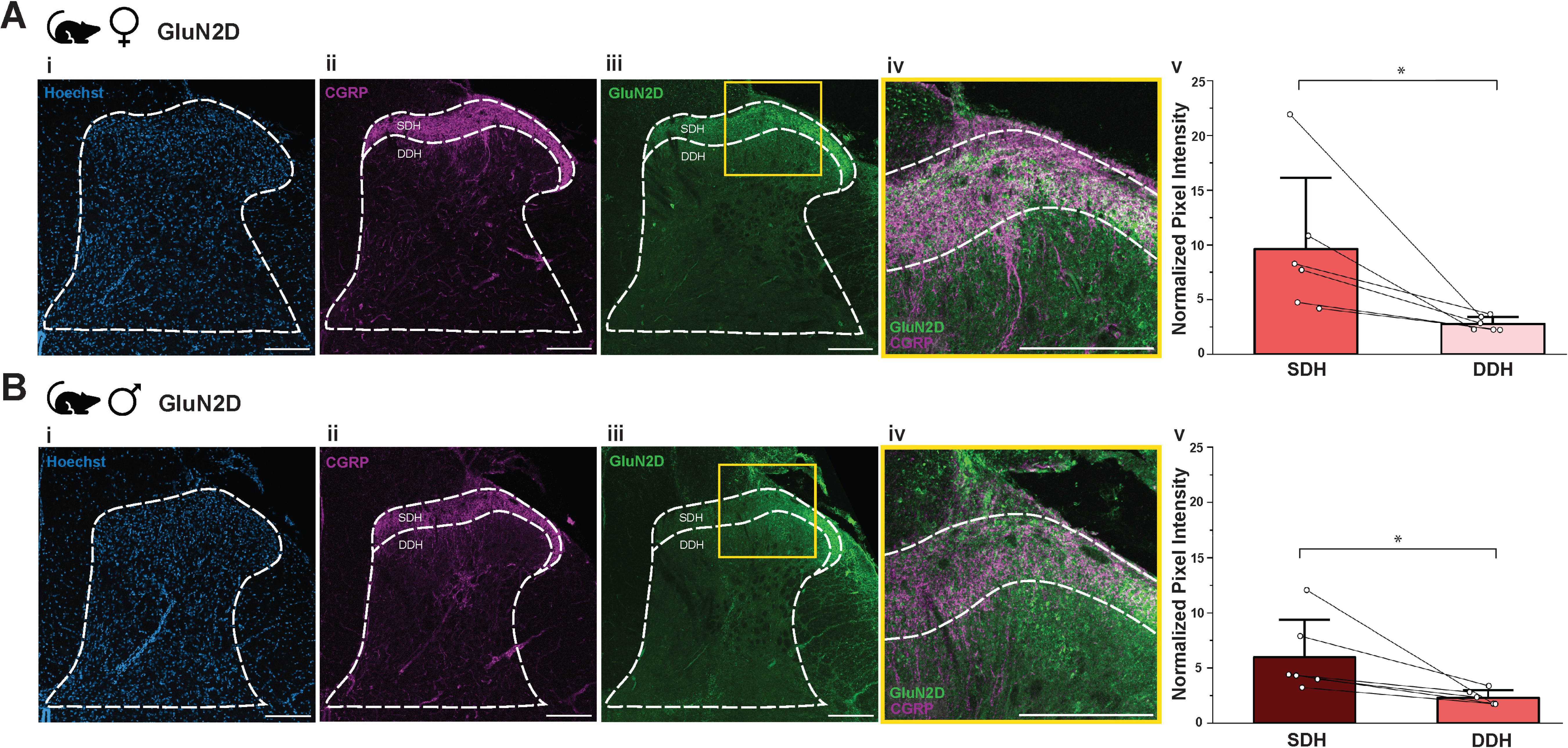
GluN2D subunits show enriched localization in the SDH compared to the DDH of the spinal dorsal horn in male and female adult rats. Representative confocal images (20× objective) from **(A)** female (L4) and **(B)** male (L5) adult rat dorsal horns showing (i) Hoechst nuclear staining to enable outlining of the dorsal horn grey matter (white dashed line) (ii) CGRP immunoreactivity marking the superficial dorsal horn (SDH; labelled, with additional marked line) as well as the remaining deeper dorsal horn (DDH), and (iii) GluN2D immunolabeling, with a yellow box highlighting the inset region shown in the panel to the right. (iv) Higher magnification images highlight GluN2D and CGRP colocalization within the SDH. Scale bars = 200 µm. (v) Quantitative analysis comparing the average expression (normalized pixel intensity) of GluN2D subunit immunoreactivity in the SDH versus the DDH of female (*n = 6* animals) and male (*n = 6* animals) adult rats aged postnatal day 90+. Data represents means ± SEM. *p<0.05.

To translate these rodent preclinical findings across species and inform potential NMDAR-targeting treatment approaches in humans, we examined the distribution of GluN2 subunits in viable spinal cord tissue from human organ donors. Using the same tissue collection, immunostaining, and analysis conditions as in rats (see Methods), we assessed GluN2 subunit localization in the SDH and DDH of spinal cord sections from female (Fig. 4-6A) and male (Fig. 4-6B) human donors. As found in rats, we observed strong immunoreactivity for both GluN2A (Fig. 4<u>(iii)</u>) and GluN2B (Fig. 5<u>(iii)</u>), with moderate levels of GluN2D (Fig. 6<u>(iii)</u>) in the dorsal horn from human donor sections. All three subunits were preferentially localized to the SDH and distribution patterns were conserved between male and female donors (Fig. 4-6<u>(iii)</u>). At a subcellular level, all three subunits were found in both synaptic neuropil as well as at cell somas, with more prominent somatic labelling in humans compared to rats (Fig. 4-6(iv)). From these initial qualitative assessments, we conclude that GluN2A, GluN2B, and GluN2D are all robustly expressed in rat and human dorsal horn cells and processes, with localization patterns that appear to be conserved across sex but that may vary from rats to humans.

**Figure 4.**
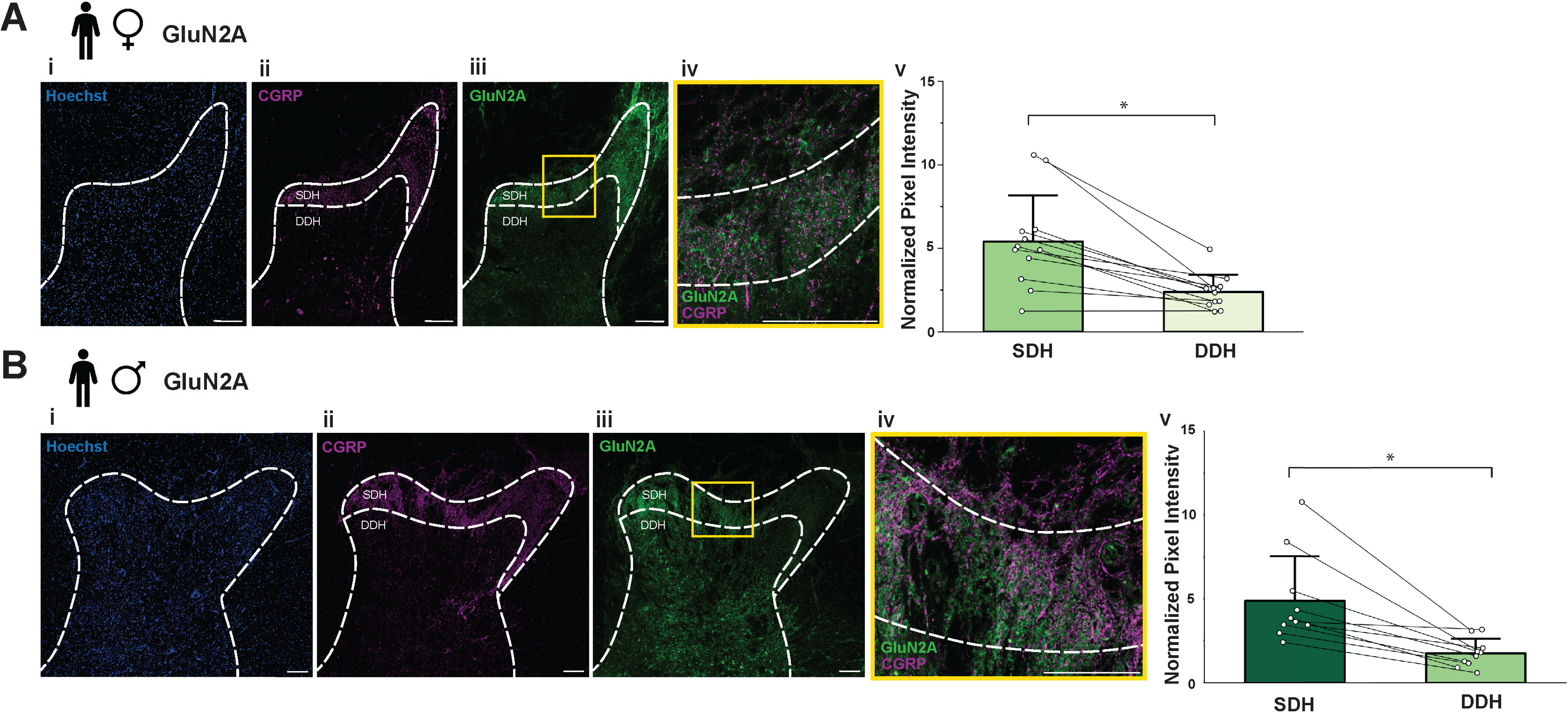
GluN2A subunits are significantly localized to the SDH compared to the DDH of the spinal dorsal horn in male and female adult humans. Representative confocal images (20× objective) from **(A)** female (L3) and **(B)** male (L5) adult human dorsal horns showing (i) Hoechst nuclear staining to enable outlining of the dorsal horn grey matter (white dashed line) (ii) CGRP immunoreactivity marking the superficial dorsal horn (SDH; labelled, with additional marked line) as well as the remaining deeper dorsal horn (DDH), and (iii) GluN2A immunolabeling, with a yellow box highlighting the inset region shown in the panel to the right. (iv) Higher magnification images highlight GluN2A and CGRP colocalization within the SDH. Scale bars = 200 µm. (v) Quantitative analysis comparing the average expression (normalized pixel intensity) of GluN2A subunits in the SDH versus the DDH of female (*n = 12* donors) and male (*n = 10* donors) adult human organ donors. Data represents means ± SEM. *p<0.05.

**Figure 5.**
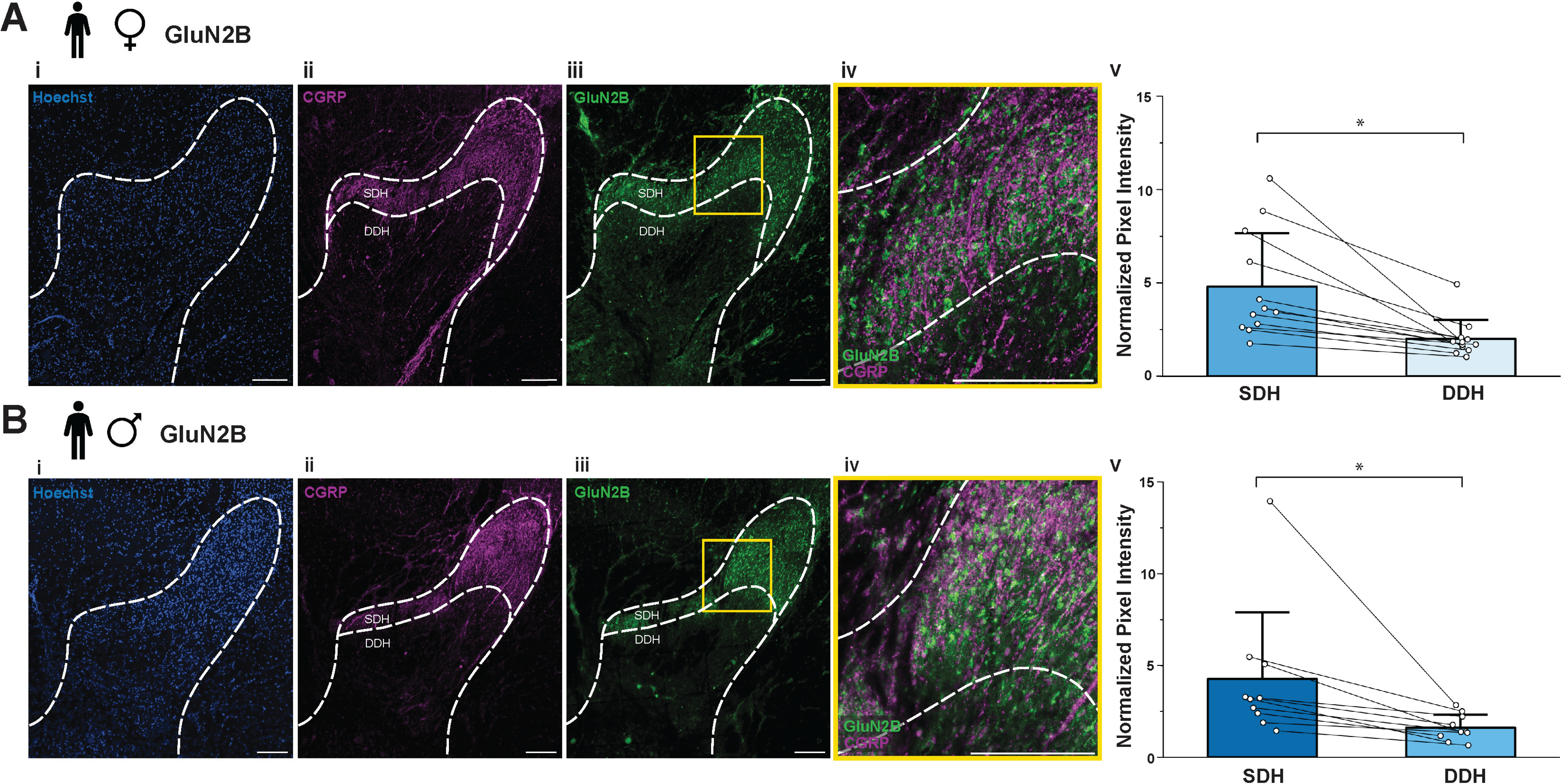
GluN2B subunits are significantly localized to the SDH compared to the DDH of the spinal dorsal horn in male and female adult humans. Representative confocal images (20× objective) from **(A)** female (L4) and **(B)** male (L3) adult human dorsal horns showing (i) Hoechst nuclear staining to enable outlining of the dorsal horn grey matter (white dashed line) (ii) CGRP immunoreactivity marking the superficial dorsal horn (SDH; labelled, with additional marked line) as well as the remaining deeper dorsal horn (DDH), and (iii) GluN2B immunolabeling, with a yellow box highlighting the inset region shown in the panel to the right. (iv) Higher magnification images highlight GluN2B and CGRP colocalization within the SDH. Scale bars = 200 µm. (v) Quantitative analysis comparing the average expression (normalized pixel intensity) of GluN2B subunits in the SDH versus the DDH of female (*n = 12* donors) and male (*n = 10* donors) adult human organ donors. Data represents means ± SEM. *p<0.05.

**Figure 6.**
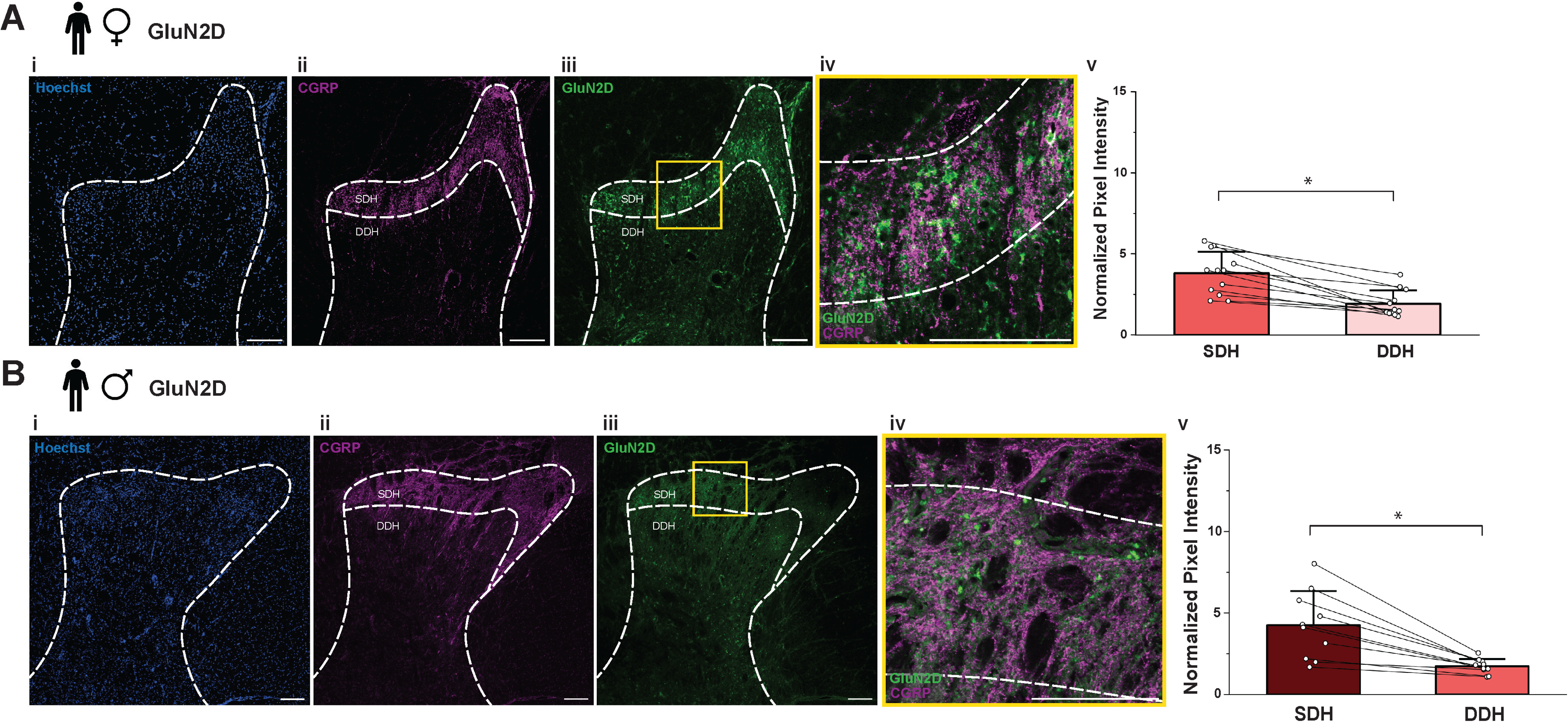
GluN2D subunits are significantly localized to the SDH compared to the DDH of the spinal dorsal horn in male and female adult humans. Representative confocal images (20× objective) from **(A)** female (L4) and **(B)** male (L5) adult human dorsal horns showing (i) Hoechst nuclear staining to enable outlining of the dorsal horn grey matter (white dashed line) (ii) CGRP immunoreactivity marking the superficial dorsal horn (SDH; labelled, with additional marked line) as well as the remaining deeper dorsal horn (DDH), and (iii) GluN2D immunolabeling, with a yellow box highlighting the inset region shown in the panel to the right. (iv) Higher magnification images highlight GluN2D and CGRP colocalization within the SDH. Scale bars = 200 µm. (v) Quantitative analysis comparing the average expression (normalized pixel intensity) of GluN2D subunits in the SDH versus the DDH of female (*n = 12* donors) and male (*n = 10* donors) adult human organ donors. Data represents means ± SEM. *p<0.05.

### Preferential Localization of GluN2A, GluN2B, and GluN2D NMDAR Subunits to the SDH is Conserved Across Sex and Species

We next quantified and statistically compared the relative total immunofluorescence (average pixel intensity per area, normalized to background) for each GluN2 subunit in the nociceptive-processing SDH (CGRP-positive) versus the somatosensory/premotor processing DDH (CGRP-negative) dorsal horn regions (Koch et al., 2018) for both species. Using this unbiased approach, we observed significantly higher immunoreactivity in the SDH compared to the DDH in female adult rats for GluN2A (Fig. 1A<u>(v)</u>; SDH = 8.02 ± 1.37; DDH = 2.95 ± 0.45; *n* = 6; *p* = 0.0033), GluN2B (Fig. 2A<u>(v)</u>; SDH = 7.13 ± 1.16; DDH = 2.21 ± 0.23; *n* = 6; *p* = 0.0046), and GluN2D (Fig. 3A<u>(v)</u>; SDH = 9.63 ± 2.66; DDH = 2.79 ± 0.26; *n* = 6; *p* = 0.042). A similar preferential localization to the SDH was also evident in the dorsal horn of adult male rats for GluN2A (Fig. 1B<u>(v)</u>; SDH = 8.54 ± 1.10; DDH = 2.91 ± 0.25; *n* = 6; *p* = 0.0029), GluN2B (Fig. 2B<u>(v)</u>; SDH = 10.59 ± 1.72; DDH = 2.48 ± 0.31; *n* = 6; *p* = 0.0026), and GluN2D subunits (Fig. 3B<u>(v)</u>; SDH = 5.99 ± 1.38; DDH = 2.32 ± 0.28; *n* = 6; *p* = 0.047). This quantification also suggests that overall GluN2 levels in the SDH increase across puberty into adulthood, as we previously used the same antibodies and staining procedures to show that GluN2A, GluN2B and GluN2D have approximately 2-3 fold higher expression in the SDH compared to background in juvenile (PD21) male and female rats (Temi et al., 2021), whereas here we find approximately 6-10 fold normalized expression for the same subunits in the SDH (Fig. 1-3(v)).

When we statistically compared the differential immunoreactivity of these GluN2 subunits between the SDH and DDH of fixed human spinal cord sections, the pattern of preferential localization to the SDH observed in rats was conserved for humans of both sexes. Specifically, for female humans, GluN2A showed significantly enhanced immunoreactivity in the SDH compared to the DDH (Fig. 4A<u>(v)</u>; SDH = 5.40 ± 0.80; DDH = 2.39 ± 0.29; *n* = 12; *p* = 0.00050), as did GluN2B (Fig. 5A<u>(v)</u>; SDH = 4.80 ± 0.83; DDH = 2.00 ± 0.29; *n* = 12; *p* = 0.0023), and GluN2D (Fig. 6A<u>(v)</u>; SDH = 3.82 ± 0.38; DDH = 1.94 ± 0.24; *n* = 12; *p* = 0.00011). Parallel differences were found in male human donors, with significantly higher levels of immunoreactivity in the SDH for GluN2A (Fig. 4B<u>(v)</u>; SDH = 4.89 ± 0.84; DDH = 1.76 ± 0.27; *n* = 10; *p* = 0.0020), GluN2B (Fig. 5B<u>(v)</u>; SDH = 4.27 ± 1.15; DDH = 1.62 ± 0.23; *n* = 10; *p* = 0.026), and GluN2D (Fig. 6B<u>(v)</u>; SDH = 4.26 ± 0.66; DDH = 1.73 ± 0.14; *n* = 10; *p* = 0.0012).

To assess whether the enhanced expression of GluN2 subunits in the SDH differs by subunit type or sex, we conducted a two-way ANOVA to compare the SDH/DDH ratio of subunit immunoreactivity in both male and female rodent (Fig. 7A<u>(i)</u>) and human (Fig. 7A<u>(ii)</u>) tissue. In rats, the analysis revealed no significant differences in the ratio among subunits (*F*(2,30) = 1.52, *p* = 0.24, *partial ^2^* = 0.09), or between sexes (*F*(1,30) = 0.401, *p* = 0.53, *partial* η = 0.013). Similarly, in human tissue, no significant subunit-related (*F*(2,60) = 0.812, *p* = 0.45, *partial* η*²* = 0.026), or sex-related (*F*(2,60) = 1.66, *p* = 0.20, *partial* η² = 0.03) differences were observed. Although we did not statistically compare across species due to differences in experimental variables such as age range, we observed that the preferential localization of GluN2B (and potentially GluN2D) to the SDH was slightly lower in humans (Fig. 7A). Altogether, these findings suggest that the enhanced localization of GluN2A, GluN2B, and GluN2D to the SDH is consistent regardless of subunit type and sex, for both rodents and humans, underscoring the critical roles of all three subunits in spinal nociceptive processing.

**Figure 7.**
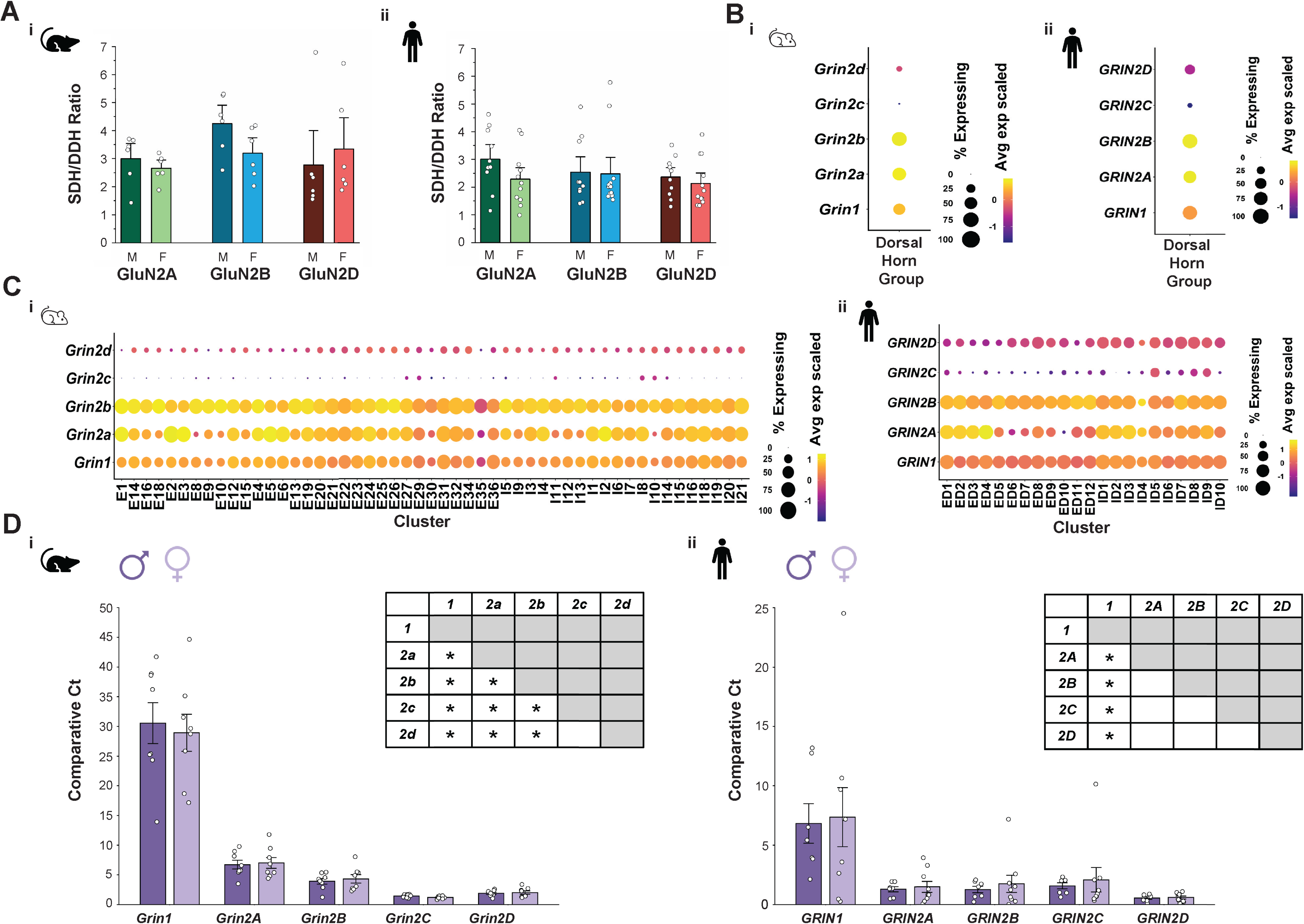
GluN2A, GluN2B, and GluN2D subunits are prominently expressed and preferentially localized to the SDH in rodent and human spinal cord of both sexes. **A)** Quantitative statistical analysis for adult (i) rats (males, *n = 6* animals; females, *n = 6* animals), and (ii) humans (male, *n = 10* donors; female, *n = 12* donors) comparing the average SDH/DDH ratio of immunoreactivity (normalized pixel intensity) for GluN2A, GluN2B, and GluN2D subunits across sex or subunits. Data represents means ± SEM. **B-C)** Dot plots showing the average expression and percent expressed for the genes encoding GluN1 (*GRIN1*), GluN2A (*GRIN2A*), GluN2B (*GRIN2B*), GluN2C (*GRIN2C*), and GluN2D (*GRIN2D*) in pooled **(B)** and individual **(C)** dorsal horn neuronal subpopulations from mouse (i) (Russ et al., 2021) and human (ii) (Yadav et al., 2023) single-cell/nucleus RNA sequencing datasets. In panel **C**, the numbering of mouse (e.g., E1, I1) and human (e.g., ED1, ID1) clusters correspond to Excit-01/Inhib-01 and Ex-Dorsal-1/Inh-Dorsal-1 clusters, respectively, as defined in the source publications (Russ et al., 2021),(Yadav et al., 2023). The average expression is represented as the color of the dot and is a *z*-score scale of log_10_(CPM +1) counts. The percent expressed is represented by the size of each dot. **D)** qRT-PCR comparative Ct analysis of dorsal horn *GRIN* gene expression in (i) rat (males, *n = 8* animals; females, *n = 8* animals) and (ii) human (males, *n = 8* donors; females, *n = 8* donors). Data are presented as mean relative expression (*2^−(^*^Δ*Ct)*^ × *1000*) normalized to the *ACTB* reference gene ± SEM, while statistical analyses were performed on ΔCt values to preserve the log-transformed scale. Pairwise comparison tables summarize post hoc comparisons between GRIN subunits, *p<0.05.

### Robust Expression of Genes Encoding GluN2A and GluN2B and Moderate Expression of GluN2D in Dorsal Horn Neuron Subpopulations of Mouse, Rat and Human Spinal Cord

To investigate the relative expression of NMDAR subunits specifically in postsynaptic dorsal horn neurons of rodents and humans, we analyzed recent single-cell/nuclei RNA sequencing datasets from mice (Russ et al., 2021) and humans (Yadav et al., 2023) for genes encoding GluN1 (*GRIN1*) and GluN2A-D (*GRIN2A-D*). In line with our immunohistochemical staining findings, pooled population analysis of all designated dorsal horn neurons demonstrated the highest expression of *GRIN2B*, followed closely by *GRIN2A* and then *GRIN1*, with moderate *GRIN2D* expression and minimal expression of *GRIN2C* in viable spinal cord tissue from both mice (Fig. 7B<u>(i))</u> and human organ donors (Fig. 7B<u>(ii))</u>.

When examining individual dorsal horn neuronal subpopulations, we identified subunit-specific differences in gene expression patterns. For the constitutive NMDAR *GRIN1* gene, the average expression and percent expressed were consistently high across all excitatory and inhibitory dorsal horn neuronal subpopulations in both mice (Fig. 7C<u>(i)</u>) and humans (Fig. 7C<u>(ii)</u>). In contrast, *GRIN2A* was highly expressed in both species but exhibited considerable heterogeneity in expression across both excitatory and inhibitory subpopulations, with high percentage and average expression in some neuronal clusters and minimal *GRIN2A* expression in others (Fig. 7C). This variability may indicate subpopulation-specific roles of GluN2A subunits in distinct aspects of somatosensory processing.

In both species, *GRIN2B* average expression was consistently higher than that of the gene encoding the obligatory GluN1 protein (*GRIN1*), with relatively uniform percent expression across cell types. Consistent with previous literature (Tölle et al., 1993; Yung, 1998; Shibata et al., 1999; Stegenga and Kalb, 2001), *GRIN2C* showed a very low overall percent expressed and average expression in the rodent and human dorsal horn (Fig. 7C). Taken together, this single-cell/nuclei dataset analysis demonstrates that the genes encoding GluN2A and GluN2B are highly expressed in mouse and human dorsal horn neurons, with moderate expression of GluN2D.

As our single-cell/nuclei sequencing analysis is restricted to post-synaptic dorsal horn neurons and is not designed nor powered for analysis by sex, we next used qRT-PCR to quantitatively compare relative subunit expression between sexes in the rat and human dorsal horn. We performed 384-well qRT-PCR on dissected rat and human dorsal horn tissue for genes encoding GluN1 (*GRIN1*) and GluN2A-D (*GRIN2A-D*) (Fig. 7D). In rats, this approach revealed a significant main effect of *Grin* gene target (*F*(4,70) = 191.26, *p* = 6.9E-37, *partial* η*^2^* = 0.92), with no significant main effect of sex (*F*(1,70) = 0.046, *p* = 0.83, *partial ^2^*= 6.5E-4) (Fig. 7D<u>(i)</u>). Bootstrapped Bonferroni-corrected post hoc analyses showed that *Grin1* expression was significantly higher than all *Grin2* subunits (*Grin2a*: [BCa 95% CI (-2.6, -1.6)], *p =* 2.2E-17 ; *Grin2b*: [BCa 95% CI (-3.5, -2.3)], *p =* 1.7E-24; *Grin2c*: [BCa 95% CI (-4.9, -4,0)], *p =* 1.9E- 35; *Grin2d* [BCa 95% CI (-4.5, -3.4)], *p =* 3.6E-32). Among *Grin2* subunits, *Grin2a* expression was significantly higher than *Grin2b* ([BCa 95% CI (-1.4, -0.21)], *p =* 3.4E-4), *Grin2c* ([BCa 95% CI (-2.7, -1.9)], *p =* 2.4E-19), and *Grin2d* ([BCa 95% CI (-2.3, -1.3)], *p =* 2.1E-14). In addition, *Grin2b* expression was significantly higher than both *Grin2c* ([BCa 95% CI (-2.0, - 1.0)], *p =* 1.6E-11) and *Grin2d* ([BCa 95% CI (-1.6, -0.4)], *p =* 2.3E-06), whereas no significant difference was observed between *Grin2c* and *Grin2d*.

In humans, there was also a significant main effect of *Grin* gene target (*F*(4,70) = 9.6, *p* = 3.0E-6, *partial* η*^2^* = 0.35), with no significant main effect for sex (*F*(1,70) = 1.0, *p* = 0.32, *partial* η = 0.01) (Fig. 7D<u>(ii)</u>). Bootstrapped Bonferroni-corrected post hoc analyses showed that *Grin1* expression was significantly higher than all *Grin2* subunits (*Grin2a*: [BCa 95% CI (-3.8, -0.65)], *p =* 0.0017; *Grin2b*: [BCa 95% CI (-4.3, -0.70)], *p =* 5.59E-4; *Grin2c*: [BCa 95% CI (-3.1, - 0.33)], *p =* 0.015; *Grin2d* [BCa 95% CI (-4.7, -2.0)], *p =* 8.9E-7). In contrast, no significant differences were observed among the *Grin*2 subunits themselves.

Overall, *Grin1* expression was higher than all *Grin*2 genes in both rats and humans, with this pattern conserved across sex, consistent with its role as the obligatory NMDAR subunit (Paoletti et al., 2013). In both species, *Grin2* subunit relative expression generally followed the expected pattern observed in our immunohistochemical and single-cell/nuclei sequencing results, with robust *Grin2a* and *Grin2b* expression and moderate *Grin2d* expression, and no significant differences in expression between males and females. However, these qRT-PCR findings suggest a potentially greater relative expression of *Grin2c* transcript in humans (Fig. 7D<u>(ii)</u>), which could be further investigated in future cell-type specific and functional studies.

### Species-Specific Asymmetry in GluN2 Subunit Distribution Across the Mediolateral Axis of the SDH

We previously demonstrated that a GluN2 subtype (GluN2B) is asymmetrically distributed across the SDH of juvenile rats in a sex-specific manner (Temi et al., 2021). To systematically study GluN2 subunit levels along the SDH mediolateral axis of adult rats in an unbiased manner, we quantified GluN2 subtype immunoreactivity across a line scan of the SDH in rat and human spinal sections (Fig. 8A). In both male and female rats, these line scans reveal a striking asymmetrical distribution across the SDH, with robust lateral enrichment of GluN2A, GluN2B and GluN2D subunits when normalized to the medial SDH region (designated at 0 μm) (Fig. 8B<u>(i-iii))</u>. A two-way ANOVA analysis revealed that there were no significant differences in the lateral/medial SDH expression ratio across subunits (*F*(2,30) = 2.45, *p* = 0.10, *partial* η*^2^* = 0.14), nor across sex (*F*(1,30) = 1.62, *p* = 0.21, *partial ^2^* = 0.05) (Fig. 8C<u>(i)),</u> indicating a robust and conserved mediolateral asymmetry of GluN2 subunit distribution in the SDH of rats.

**Figure 8.**
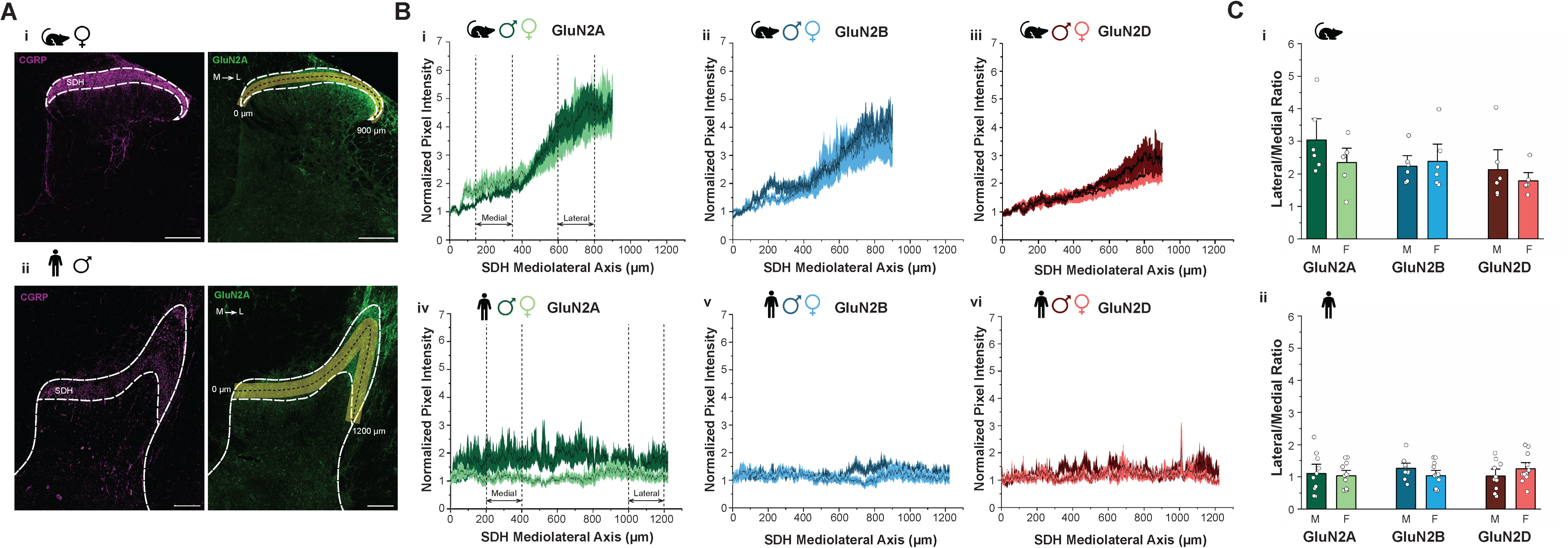
GluN2A, GluN2B, and GluN2D subunits are asymmetrically distributed along the mediolateral SDH axis of male and female rat, but not human, spinal cord. **A)** Representative confocal images (20x objective) of CGRP (*red*) and GluN2A (*green*) immunohistochemical staining in rat (i) and human (ii) dorsal horns with line scans (*in yellow*) that were used for immunoreactive quantification drawn across the mediolateral axis of the SDH (rats: 0 µm = medial start point, 900 µm = lateral end point; humans: 0 µm = medial start point, 1200 µm = lateral end point). Scale bars = 200 µm. B) *Top*, average mediolateral line scan of GluN2A (i), GluN2B (ii) and GluN2D (iii) subunit expression (normalized pixel intensity) across the SDH in male (*n = 6* animals) and female (*n = 6* animals) rats. *Bottom*, average mediolateral line scan for GluN2A (iv), GluN2B (v) and GluN2D (vi) subunit expression across the SDH in male (*n = 10* donors) and female (*n = 12* donors) humans. Line scans were normalized by dividing all pixel intensity measurements per area along the line by the average pixel intensity within a defined medial reference range (50-200 μm). Data represents mean (*black line*) ± SEM. (*coloured shading*). Dashed black lines indicate the medial and lateral regions used for ratio analysis (rats: medial = 200-400 µm, lateral = 600-800 µm; humans: medial = 200-400 µm, lateral = 1000-1200 µm). **C)** Quantitative statistical analysis for adult rats (i; males, *n = 6* animals; females, *n = 6* animals), and humans (ii; male, *n = 10* donors; female, *n = 12* donors) comparing the average lateral to medial SDH ratio for GluN2A, GluN2B, and GluN2D subunits across sex or subunits. Data represents means ± SEM.

In contrast, humans displayed a generally uniform mediolateral distribution of GluN2 subunits across the SDH. Line scans revealed stable immunoreactivity levels across the human SDH mediolateral axis for GluN2A (Fig. 8B<u>(iv)),</u> GluN2B (Fig. 8B<u>(v))</u> and GluN2D (Fig. 8B<u>(vi))</u> subunits. Furthermore, no significant differences in the lateral/medial SDH ratio were found across subunits (*F*(2,60) = 1.94, *p* = 0.15, *partial ^2^* = 0.06) or sexes (*F*(1,60) = 0.662, *p* = 0.42, *partial* η*^2^* = 0.011) in humans (Fig. 8C<u>(ii)).</u> Therefore, while rats exhibit a prominent lateral enrichment of GluN2 subunits within the SDH, suggesting potential region-specific differences in synaptic properties, this asymmetry is almost entirely absent in humans, underscoring a key species difference in NMDAR localization across the SDH.

### Increased Expression of GluN2A Protein and Transcript But Not Primary Afferent or Neuronal Markers Across the Mediolateral SDH Axis of Adult Rats

We next investigated whether the heightened localization of GluN2 NMDAR subunits to the lateral SDH could be driven by differences in levels of presynaptic innervation or postsynaptic neuronal density across the rodent SDH mediolateral axis. The neuropeptide CGRP is primarily localized within SDH presynaptic terminals of innervating peptidergic primary afferent fibres (Gibson et al., 1984; Chung et al., 1988). We examined CGRP localization across the SDH mediolateral axis in female rats using the same analysis methods as for GluN2 subunits (Fig. 9A), given that no sexes differences were observed in the preceding findings. Line scan analysis of CGRP immunoreactivity revealed a strikingly uniform distribution across the SDH mediolateral axis (Fig. 9A<u>(ii))</u>, with no significant difference in CGRP levels between the medial and lateral SDH regions (Supplementary Fig. 2B, medial SDH = 1.07 ± 0.06; lateral SDH = 1.01 ± 0.11; *n* = 6; *p* = 0.49). These findings suggest that the preferential localization of GluN2 subunits to the lateral SDH of adult rats is not driven by variations in presynaptic afferent terminal density.

**Figure 9.**
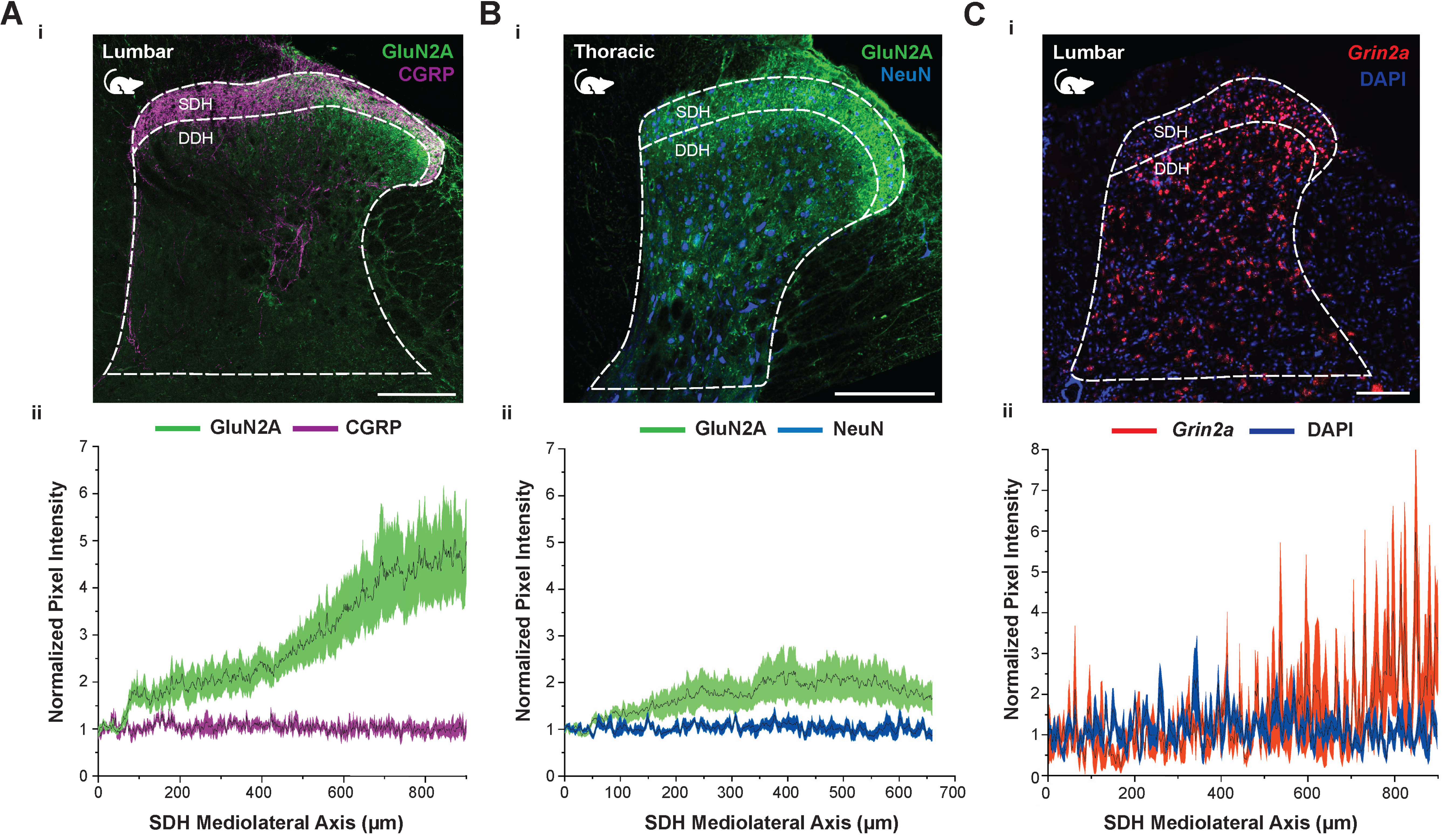
GluN2A protein and transcript, but not primary afferent and neuronal markers, are enriched in the lateral SDH of adult rats. rats. **A)** Expression of presynaptic CGRP afferents does not correlate with the asymmetrical distribution of GluN2A subunits across the SDH in adult rats. (i) Representative confocal image (20x objective) of GluN2A (*green*) and CGRP (*magenta*) immunohistochemical staining in the lumbar dorsal horn of a female adult rat (PD90+). (ii) Average mediolateral line scan of normalized GluN2A and CGRP expression (normalized pixel intensity) across the SDH (0 µm = medial start point) in female adult rats (*n = 6* animals). **B)** Expression of GluN2A subunits, but not a neuronal marker (NeuN) is preferentially localized to the lateral SDH in adult rat thoracic spinal cord. (i) Representative confocal image (20x objective) of GluN2A (*green*) immunohistochemical staining co-labeled with NeuN neuronal marker (*blue*) in the thoracic dorsal horn of a female adult rat (PD90+). (ii) Average mediolateral line scan of normalized GluN2A and NeuN expression (normalized pixel intensity) across the thoracic SDH (0 µm = medial start point) in adult rats (*n = 6* animals; 3 male, 3 female). **C)** GluN2A RNA expression aligns with the clear enhancement of GluN2A protein expression in the lateral SDH of adult rats. (i) Representative RNAscope image showing the co-labeling of mRNA expression for *Grin2a* (*red, n = 3* animals) and nuclei counterstained with DAPI (*blue, n =* 5 animals) in the lumbar dorsal horn of a female adult rat (PD90+). (ii) Average mediolateral line scan of normalized *Grin2a* and DAPI expression (normalized pixel intensity) across the SDH (0 µm = medial start point). Scale bars = 200 µm. Data represents mean (*black line*) ± SEM. (*coloured shading*).

To test if asymmetrical GluN2 distributions are conserved across spinal cord regions and potentially influenced by variations in neuronal density, we examined thoracic adult rat spinal cord tissue of both sexes, with co-staining for GluN2A alongside a neuronal marker, NeuN (Mullen et al., 1992) (Fig. 9B). Line scan analysis revealed a consistent, albeit less pronounced, increase in GluN2A immunostaining in the thoracic lateral SDH, while NeuN immunoreactivity remained uniformly distributed across the mediolateral axis (Fig. 9B<u>(ii)).</u> We found that GluN2A levels significantly increased in the lateral SDH compared to the medial SDH (Supplementary Fig. 3A, medial SDH = 1.48 ± 0.19; lateral SDH = 2.07 ± 0.39; *n* = 6; *p* = 0.039), while NeuN protein levels did not differ across the SDH mediolateral axis (Supplementary Fig. 3B, medial SDH = 1.03 ± 0.07; lateral SDH = 0.99 ± 0.12; *n* = 6; *p* = 0.68). Thus, the asymmetrical distribution of GluN2A is maintained across multiple rat spinal cord regions and is not attributable to differences in neuronal density across the SDH mediolateral axis.

The differences in GluN2 subunit protein abundance between the medial and lateral SDH could be driven by differences in gene expression and/or protein translation, trafficking and degradation. An RNAscope *in situ* hybridization assay was therefore performed using an mRNA probe for the rat GluN2A subunit gene (*Grin2a*) in female lumbar spinal cord tissue to test whether the GluN2A transcript expression pattern corresponds with the asymmetry in protein distribution (Fig. 9C<u>(i)</u>). Line scan analyses across the SDH revealed that *Grin2a* mRNA levels increased from the medial to the lateral SDH, as observed for GluN2A protein (Fig. 8B<u>(i)</u>), while nuclei counterstained with DAPI displayed a uniform distribution across the SDH mediolateral axis ((Fig. 9C<u>(ii)</u>). These findings further validate the observed asymmetry in GluN2 expression across the SDH and suggest that *Grin2a* RNA expression closely parallels the enhanced presence of GluN2A protein in the lateral SDH.

### The Abundance and Lateral Localization of the GluN2A Subunit in the SDH Increases Over Late Postnatal Development in Male and Female Rats

In contrast to the striking preferential localization of GluN2A to the lateral SDH of adult rats observed here (Fig. 7A<u>(i)</u>; Fig. 8B<u>(i)</u>), we have previously found that GluN2A subunits are evenly distributed at low levels across the dorsal horn of juvenile male and female rats (Temi et al., 2021). To directly investigate potential changes in GluN2A localization across later postnatal development, we replicated our immunohistostaining of GluN2A in juvenile (PD21) male and female rat tissue while, in parallel, we also stained post-pubertal adolescent (PD42) tissue of both sexes. The distribution of GluN2A immunoreactivity was then analyzed at these PD21 and PD42 developmental stages and compared to our staining results from adult (PD90+) rat tissue (Fig. 10A).

**Figure 10.**
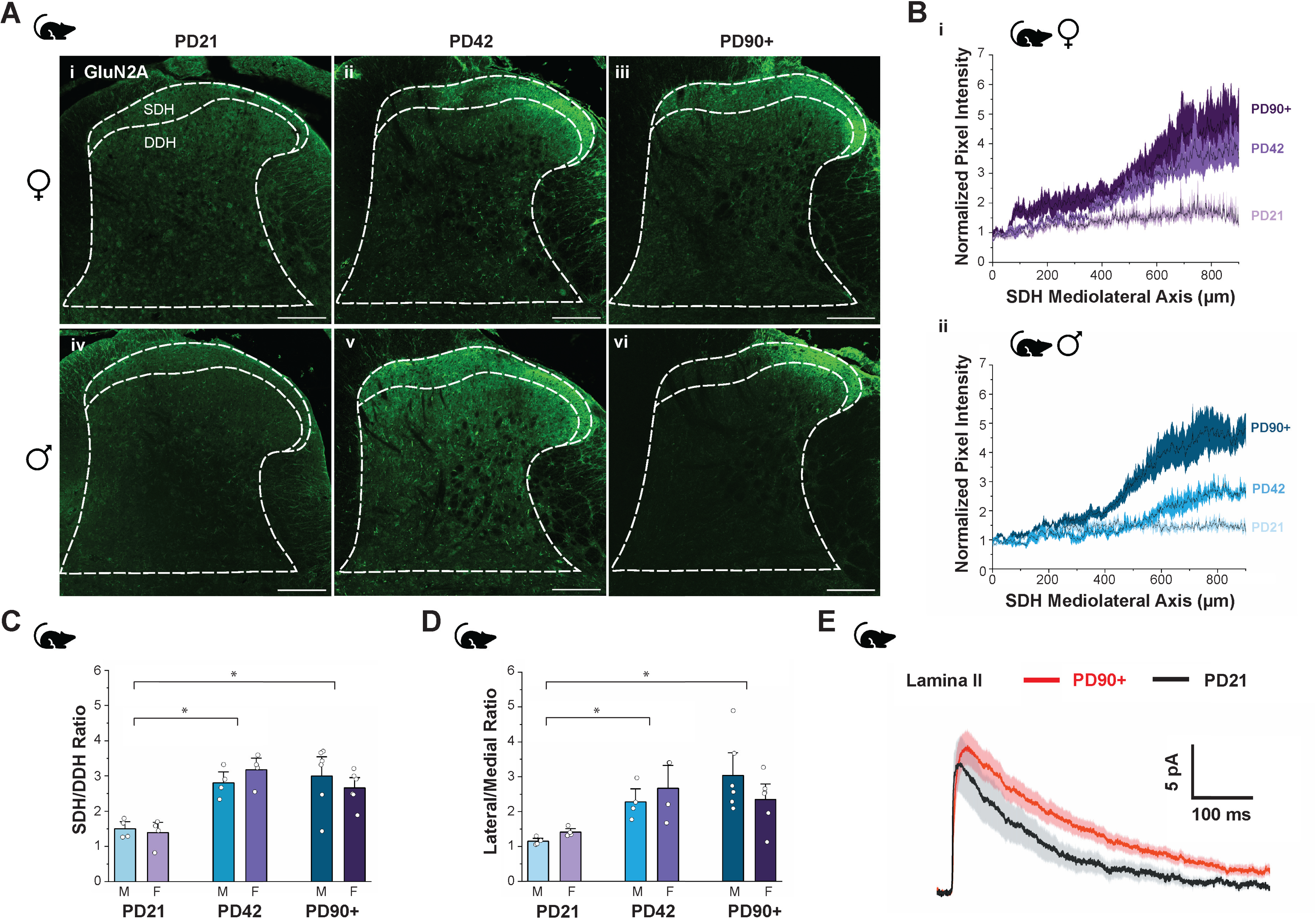
GluN2A subunit expression and lateral SDH localization increases across adolescent development in male and female rats. **A)** Representative confocal images (20× objective) of GluN2A subunit immunostaining (*in green*) in the spinal dorsal horn of female (*top*) and male (*bottom*) rats aged postnatal day 21 (i, iv; juvenile), postnatal day 42 (ii, v; adolescent), and postnatal day 90+ (iii, vi; adult). Scale bars = 200 µm. **B)** Average mediolateral line scan of normalized GluN2A subunit expression (normalized pixel intensity) across the SDH in female rats (i; *n = 4*) and male rats (ii; *n = 4* animals) demonstrating a transition in GluN2A localization across three developmental stages. Data represents mean (*black line*) ± SEM. (*coloured shading*). **D-E)** Quantitative statistical analysis comparing the SDH/DDH ratio (**C**) and the lateral SDH/medial SDH ratio (**D**) for GluN2A subunits between sexes (males: *blue*, *n = 4* animals; females: *purple, n = 4* animals) and across various developmental timepoints (PD21, PD42, PD90+). Data represents means ± SEM. *p<0.05. **E)** Averaged +60 mV mEPSC traces in lamina II SDH spinal cord neurons of juvenile rats (PD20 to PD22; *black*; *n =* 5 neurons from 3 animals) compared to adult rats (PD90+; *red; n =* 11 neurons from 6 animals). Scale bars = 100 ms (*x-axis*); 5pA (*y-axis*). Data represents means ± SEM.

Similar to our past findings (Temi et al., 2021), we found no significant difference between GluN2A immunoreactivity in the SDH versus the DDH at PD21, confirming relatively uniform levels of GluN2A protein across the juvenile dorsal horn in both sexes. Specifically, SDH and DDH normalized pixel intensity values were comparable in females (Supplementary Fig. 4A<u>(i);</u> SDH = 2.75 ± 0.70; DDH = 1.96 ± 0.31; *n* = 4; *p* = 0.19), and males (Supplementary Fig. 4A<u>(ii);</u> SDH = 2.90 ± 0.62; DDH = 1.92 ± 0.39; *n* = 4; *p* = 0.052). However, by PD42, a dramatic shift emerged, with enhanced levels of GluN2A protein in the nociceptive-processing SDH for both females (Supplementary Fig. 4B<u>(i);</u> SDH = 8.35 ± 1.61; DDH = 2.66 ± 0.50; *n* = 4; *p* = 0.016) and males (Supplementary Fig. 4B<u>(ii);</u> SDH = 6.38 ± 0.72; DDH = 2.27 ± 0.16; *n* = 4; *p* = 0.0070).

We also assessed the localization of the GluN2A subunit across the mediolateral axis of the SDH across this later period of postnatal development for both sexes. Normalized line scans at PD21 revealed relatively uniform GluN2A across the mediolateral axis of the SDH in female (Fig. 10B<u>(i)</u>) and male (Fig. 10B<u>(ii)</u>) rats. However, quantitative analysis combining sexes revealed a modest but significant increase in GluN2A in the lateral compared to medial SDH (Supplementary Fig. 5A, medial SDH = 1.25 ± 0.06; lateral SDH = 1.59 ± 0.09; *n* = 8; *p* = 0.0035). In contrast, a dramatic shift in GluN2A localization emerged post-puberty. Normalized line scans for late adolescent rats (PD42) demonstrated a pronounced lateral enrichment in both females and males (Fig. 10B), which was supported by quantitative analysis (Supplementary Fig. 5B, medial SDH = 1.24 ± 0.07; lateral SDH = 3.02 ± 0.31; *n* = 8; *p* = 0.00060). This lateral predominance became even more evident in adulthood (Fig. 10B), collectively indicating a substantial post-pubertal reorganization of GluN2A localization within the SDH, potentially reflecting the maturation of somatotopically-organized circuitry along the mediolateral axis.

To statistically compare the effects of age and sex on GluN2A levels across development, a two-way ANOVA was conducted to evaluate the SDH/DDH ratio of pixel intensity across these factors (Fig. 10C). A significant main effect of age was observed (*F*(2,22) = 19.4, *p* = 1.4E-5, *partial ^2^* = 0.6), with no significant main effect for sex (*F*(1,22) = 0.017, *p* = 0.90, *partial ^2^* = 0.001). Bootstrapped Bonferroni-corrected post hoc analyses revealed a significantly lower SDH/DDH ratio in PD21 rats compared to both PD42 rats ([BCa 95% CI (-1.9, -1.2)], *p* = 4.4E-5), and PD90+ rats ([BCa 95% CI (-1.1, -0.96)], *p* = 5.6E-5). However, no significant difference was observed between PD42 and PD90+ rats ([BCa 95% CI (-0.30, 0.68)], *p* = 1.0). These results validate an increase in the SDH/DDH ratio of GluN2A across developmental stages, plateauing between late adolescence and adulthood, with no differences attributable to sex.

Similarly, a two-way ANOVA was performed to analyze the lateral/medial SDH expression ratio for GluN2A across juvenile, adolescent, and adult age points (Fig. 10D). A significant main effect of age was found (*F*(2,22) = 9.88, *p* = 0.00087, *partial* η*^2^* = 0.47), but no significant main effect for sex (*F*(1,22) = 0.002, *p* = 0.97, *partial ^2^*= 0.000). Bootstrapped Bonferroni-corrected post hoc analyses confirmed that the lateral/medial SDH ratio was significantly higher in both PD42 ([BCa 95% CI (0.72, 1.7)], *p* = 9.6E-3) and PD90+ ([BCa 95% CI (0.87, 2.0)], *p* = 8.8E-4) rats compared to the PD21 rats. However, no significant difference was found between PD42 and PD90+ ([BCa 95% CI (-0.88, 0.43)], *p* = 1.0). These findings confirm a developmental stage-dependent shift in the asymmetrical distribution of GluN2A expression, with notable increases in the lateral/medial SDH ratio beginning post-puberty and continuing into adulthood, without any significant sex-based differences.

To investigate the functional implications of the observed variations in GluN2 subunit SDH levels across late postnatal development, we recorded outward NMDAR-dominated miniature excitatory postsynaptic currents (mEPSCs) at +60 mV in lamina II_outer_ neurons (Dedek and Hildebrand, 2024) from juvenile (PD20-PD22) and adult (PD90+) rats. We previously found that the charge transfer, decay constants and relative contributions of GluN2A versus GluN2B subunits for NMDAR mEPSCs in lamina II neurons does not change between PD7 and PD21 (Mahmoud et al., 2020), in contrast to the GluN2B to GluN2A early postnatal developmental switch that occurs in most brain neurons (Paoletti et al., 2013). Surprisingly, we discovered here that outward mEPSCs in adult (PD90+) lamina II_outer_ neurons exhibited larger amplitudes and greater associated charge transfers as well as slower decay kinetics compared to those in juvenile (PD20-22) male rats (Fig. 10E). Specifically, the average charge transfer from 40 ms to 500 ms after mEPSC onset was 2.50 ± 0.22 pC (*n* = 11) for adult lamina II neurons, which was nearly double that observed in juveniles (1.31 ± 0.39 pC, *n* = 5, *p* = 0.011). Furthermore, the decay constant (τ_decay_) was longer in adults (198.1 ± 11.1 ms) compared to juveniles (128.1 ± 15.9 ms). In combination with our above-reported increase in GluN2A, GluN2B, and GluN2D SDH levels in adult compared to juvenile rats, these results demonstrate an increase in GluN2 subunit expression and associated functional synaptic responses within the SDH across late postnatal development, from juvenile to adult age points.

## DISCUSSION

We report here that the GluN2A and GluN2B NMDAR subunits are highly expressed at both transcript and protein levels in the dorsal horn of rat and human spinal cord, with moderate expression of GluN2D. All three of these GluN2 subunits are preferentially localized to the SDH and have distribution patterns that are conserved between sexes. In comparing results across species, we found subtle differences in the relative expression between subunits, but a striking difference where all three GluN2 subunits are enriched in the lateral SDH for rats but not humans. Moreover, we discovered a dramatic reorganization of GluN2 expression in the dorsal horn across rodent late postnatal development. For the GluN2A subunit, which has a robust degree of heightened lateral SDH expression in adult rats, we observed diffuse expression in juvenile (PD21) rats, with a large increase in immunoreactivity in the SDH and enrichment across the mediolateral axis in adolescent male and female rats (PD42) that continues into adulthood. This increase in SDH NMDAR subunits across late development is also reflected at a functional level, as NMDAR-mediated synaptic responses in lamina II neurons are larger in adult rats compared to juvenile rats.

The combination of our immunohistochemical and single-cell/nuclei sequencing results suggests species differences in NMDAR subunit expression, with potentially less pronounced expression of GluN2B and more expression of GluN2D in the dorsal horn of humans compared to rodents (Fig. 7A-C). This aligns with what we have observed at a functional level for NMDAR synaptic responses in human vs rat SDH neurons. In patch-clamp recordings on viable lamina I neurons from rat and human organ donor tissue, we found that GluN2B NMDARs mediate a smaller component of NMDAR mEPSCs in humans versus rats (Dedek et al., 2024). Moreover, a rare subset of synaptic NMDAR events in human lamina I neurons had a large amplitude and slow GluN2D-like decay constants, which were not observed in rats (Dedek et al., 2024). The potential reduced role for GluN2B NMDARs in human spinal pain processing neurons could partly explain why GluN2B-targetting therapeutics have shown efficacy in rodent preclinical pain models but have failed in human clinical trials (Liu et al., 2020), in combination with the differential dysregulation of GluN2B NMDARs between sexes (Dedek et al., 2022). In terms of promising new directions, given that GluN2D NMDAR subunits are not prominent at mature brain synapses (Paoletti et al., 2013) and yet have robust expression and functional contributions in SDH neurons from rodents to humans, GluN2D may represent a promising potential therapeutic target for pain. Notably, while qRT-PCR analysis did not reveal increased *GRIN2D* expression in humans, it suggested a potential greater relative expression of *GRIN*2C compared to rats (Fig. 7D(ii)). Accordingly, these findings raise the possibility that GluN2C, alongside GluN2D, may contribute to prolonged synaptic NMDAR responses in human dorsal horn nociceptive circuits (Sundström et al., 1997).

One caveat when comparing and reconciling our complementary immunostaining data with single-cell/nuclei RNA sequencing and qRT-PCR results is the source(s) of transcript/protein. While the latter transcript-based approaches will reflect expression that is dominated by dorsal horn cells (and specifically postsynaptic dorsal horn neurons for our single-cell/nuclei results), immunostaining will label GluN2 protein at a host of cellular and subcellular locations. This can include localization of these NMDAR subunits to non-neuronal cell types as well as presynaptic terminals of both primary afferents and descending efferents (Deng et al., 2019; Verkhratsky and Chvátal, 2020; Dedek and Hildebrand, 2022). Thus, the roles of these detected GluN2A, GluN2B, and GluN2D subunit proteins in dorsal horn nociceptive circuitry is not necessarily limited to postsynaptic dorsal horn responses. We also note that although the GluN2 antibodies that we used here have been previously optimized for immunohistochemistry by our group and others (Temi et al., 2021) and that our staining results align with our other experimental evidence as well as past spinal cord staining (Abe et al., 2005; Hummel et al., 2008; Ji et al., 2015), we cannot exclude that some GluN2 immunoreactivity in our rat and human tissue may be driven by non-specific binding.

Sex is a critical biological variable in biomedical research and has profound implications for the understanding and treatment of pain (Mogil, 2020; Shansky and Murphy, 2021). We found here that the expression and distribution of distinct GluN2 subunits – including GluN2A, GluN2B, and GluN2D – within the pain-processing SDH is conserved across sex for both rodents and humans. Similarly, we have also reported that the relative contributions of GluN2 subunits towards rodent and human synaptic NMDARs (Dedek et al., 2024) as well as their general expression pattern in juvenile rats (Temi et al., 2021) are also concerned across sex in pain-naïve states. This is in contrast to the discovery of sexually divergent mechanisms that dysregulate spinal NMDARs in rodent and human tissue preclinical models of pathological pain (Dedek et al., 2022; Hayashi et al., 2024). From these combined findings we therefore propose that the molecular and functional properties of SDH NMDARs are conserved across sex in baseline, physiological states, but that there are profound and clinically relevant sex differences in how specific spinal NMDAR subtypes are dysregulated to drive chronic pain.

A surprising twist in this study was the finding that GluN2A, GluN2B, and GluN2D subunits are all preferentially localized to the lateral half of the SDH in adult rats but not in humans. Although this enrichment of GluN2 expression in the lateral SDH has not been explicitly investigated in past rodent immunohistochemical studies, revisiting staining patterns with an eye on mediolateral distribution validates that both GluN2D (Hummel et al., 2008) and GluN2B (Abe et al., 2005; Ji et al., 2015) subunits are enriched in the lateral SDH of adult rodents. This heightened lateral expression of GluN2 subunits is not a general phenomenon driven by gross anatomical differences, as levels of presynaptic afferents, neuronal density, and another type of synaptic receptor (CB1 (Parnell et al., 2023)) are all evenly distributed across the SDH mediolateral axis. In terms of physiological implications, the mediolateral axis represents a somatotopic map where the medial dorsal horn receives distal input from the limbs and digits and the lateral portion receives proximal projections from the trunk of the body in rats (Brown et al., 1997), mice (Odagaki et al., 2019), and cats (Wilson and Snow, 1988). Recent work identifying molecular gradients that organize somatotopic mapping in the dorsal horn further supports the idea that spatially distinct afferent input streams are organized along the mediolateral axis (Sangster et al., 2026), potentially giving rise to region-specific synaptic properties. Interestingly, we found less of a lateral increase for GluN2A in the SDH of thoracic versus lumbar rat spinal cord, which may be because mediolateral somatotopic gradients vary across the rostro-caudal axis of rats (Brown et al., 1997). This observed lateral enrichment of GluN2 NMDAR subunits suggests that the physiological roles for NMDARs and their associated mechanisms of plasticity may vary by body region and type of incoming afferents, with potentially enhanced roles for NMDARs in regulating SDH excitability for proximal body regions, and hairy versus glabrous skin (Dickie et al., 2019; Hoheisel et al., 2025). Thus, neuronal location across the mediolateral axis is an important variable to be tracked in future investigations of SDH excitability and plasticity, with follow-up studies investigating how these molecular determinants of spinal cord somatotopy are potentially altered by pathological pain states in both rodents and humans.

Unlike the early postnatal changes in NMDAR GluN2 expression linked to activity-dependent plasticity and circuit wiring of the brain (Paoletti et al., 2013), we found here that GluN2A distribution in the dorsal horn changes dramatically across later postnatal development for male and female rats. Past research on the development of dorsal horn nociceptive circuits has focused on the striking changes in connectivity and plasticity between peripheral afferents and postsynaptic dorsal horn neurons that lead to the development of peripheral receptive fields in early postnatal stages (PD3-PD21) (Fitzgerald and Jennings, 1999; Koch and Fitzgerald, 2013). However, refinement of this afferent fiber-dorsal horn connectivity is an NMDAR-dependent process that may continue beyond juvenile states, with evidence for potential NMDAR-driven changes into adulthood (Beggs et al., 2002; Granmo et al., 2008). Thus, the striking changes in GluN2A localization from juvenile stages through puberty into adulthood may suggest activity-dependent processes are prominent in shaping and refining SDH nociceptive circuitry throughout the lifespan for both males and females, including across the somatotopically-organized mediolateral axis of the SDH. This molecular refinement appears to be dependent on sexual maturation, as the enrichment of GluN2A to the lateral SDH was closer to final adult rat values at PD42 for females (Fig. 10B(i)) compared to males (Fig. 10B(ii)) and puberty is complete at this timepoint for females but not males (Bell, 2018). Given the increased prevalence of chronic pain across the lifespan (Domenichiello and Ramsden, 2019), understanding these unique molecular changes in SDH nociceptive processing that are initiated during puberty and continue into adulthood may help uncover novel approaches for the treatment of pain.

## ACKNOWLEDGEMENTS

We are grateful to organ donors and their families for their extremely generous gift; this study would not have been possible without them. We thank the intensive care unit and operating room staff at the Ottawa Hospital, Civic Campus for donating their time and energy to make the human tissue collection in this study possible. Thank you to the research staff and coordinators Suzan Chen, Lei Zhou, Angela Auriat, Jessica Parnell, Harleen Kaur, and Maitreya Patel. This research was supported in part by the Intramural Research Program (ZIA NS003153 to AJL) of the National Institutes of Health (NIH), National Institute of Neurological Disorders and Stroke (NINDS). The contributions of the NIH author(s) were made as part of their official duties as NIH federal employees, are in compliance with agency policy requirements, and are considered Works of the United States Government. However, the findings and conclusions presented in this paper are those of the author(s) and do not necessarily reflect the views of the NIH or the U.S. Department of Health and Human Services.

## Data Availability

The data that support the findings of this study are available from the corresponding author, upon reasonable request.

