## Supplementary Information for "Expression and localization of NMDA receptor GluN2 subunits in dorsal horn pain circuits across sex, species, and late postnatal development"

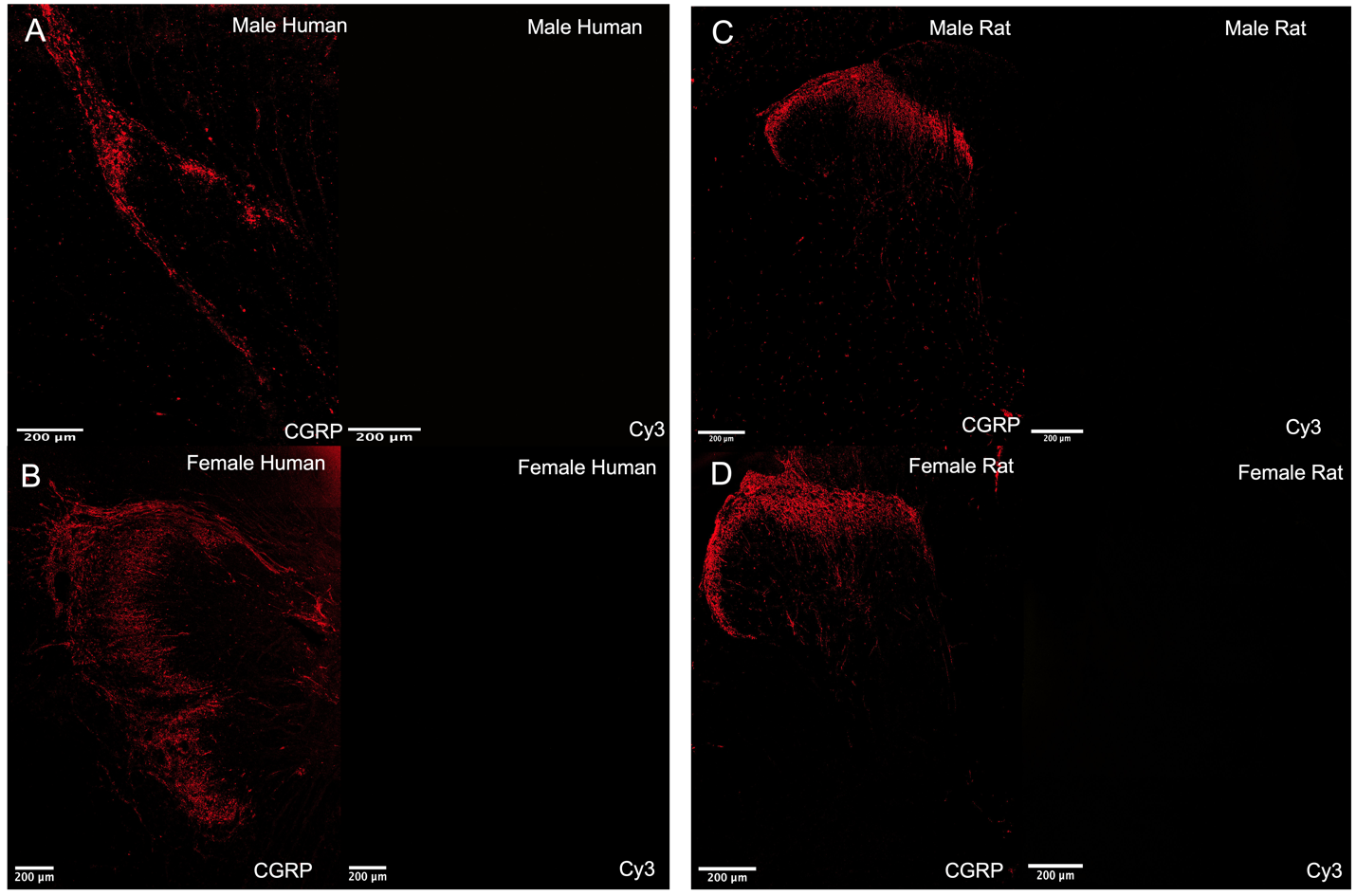

**Supplementary Figure 1. Immunohistochemical no primary GluN2 antibody controls in adult rats and humans of both sexes.** Representative confocal images (20x objective) of **(A)** male human donor, **(B)** female human donor, **(C)** male rat, and **(D)** female rat samples stained with CGRP (*left*) and no evident GluN2 primary antibody staining (*right*), indicating proper binding of secondary antibodies. Scale bars = 200 µm.

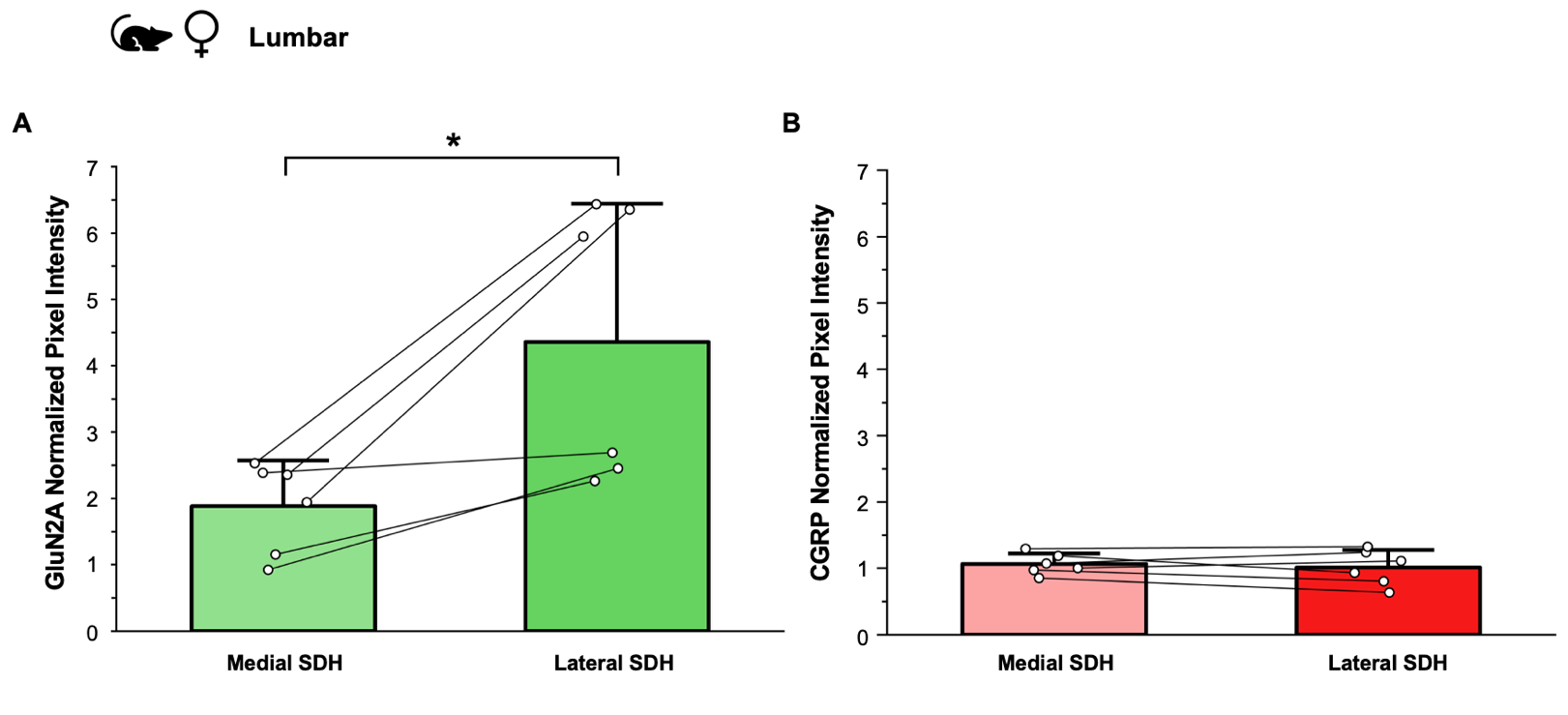

**Supplementary Figure 2.** **GluN2A subunit expression, but not presynaptic CGRP, is significantly increased in the lateral SDH compared to the medial SDH in the adult rodent spinal dorsal horn.** Quantitative analysis in adult female rats (*n* = 6 animals) comparing the mean expression (normalized pixel intensity) for **(A)** GluN2A and **(B)** CGRP in the medial SDH versus the lateral SDH of lumbar spinal cord tissue. Data represents means ± SEM. *p<0.05.

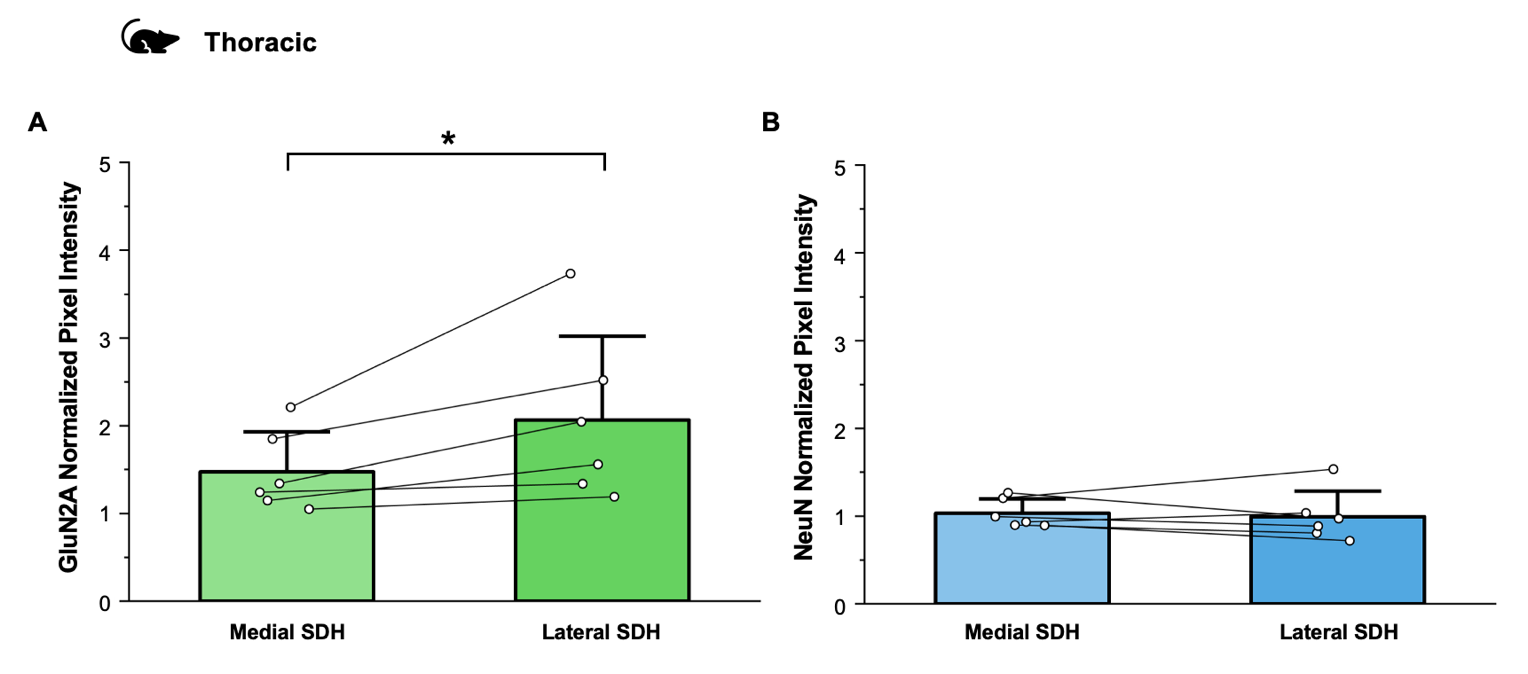

**Supplementary Figure 3.** **GluN2A subunits, but not the neuronal marker NeuN, exhibit preferential localization to the lateral SDH, a pattern conserved in the adult rat thoracic spinal cord.** Quantitative analysis of thoracic spinal cord tissue from adult rats (*n* = 6 animals [3 females, 3 males) and comparing the mean expression (normalized pixel intensity) for **(A)** GluN2A, **(B)** NeuN between the medial and lateral SDH, with data combined across sexes. Data represents means ± SEM. *p<0.05.

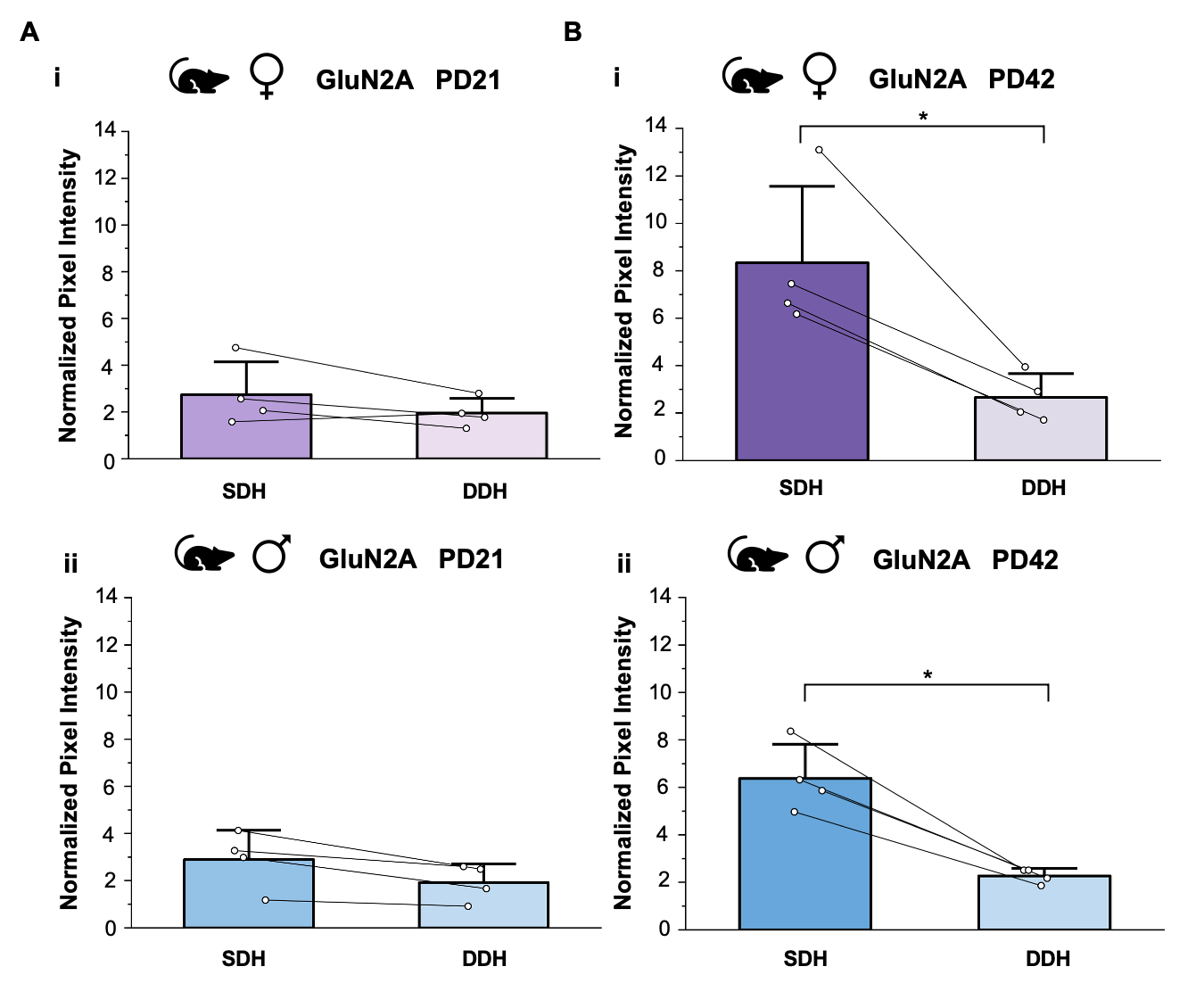

**Supplementary Figure 4. GluN2A subunit expression is significantly localized to the SDH compared to the DDH in adolescents but not in juveniles of both sexes.** Quantitative analysis comparing the average expression of GluN2A subunits (normalized pixel intensity) in the SDH versus the DDH of **(A)** juvenile rats aged postnatal day 21 and **(B)** adolescent rats aged postnatal day 42 for both sexes [**(i)** female, *n* = 4 animals; **(ii)** male, *n* = 4 animals]. Data represents means ± SEM. *p<0.05.

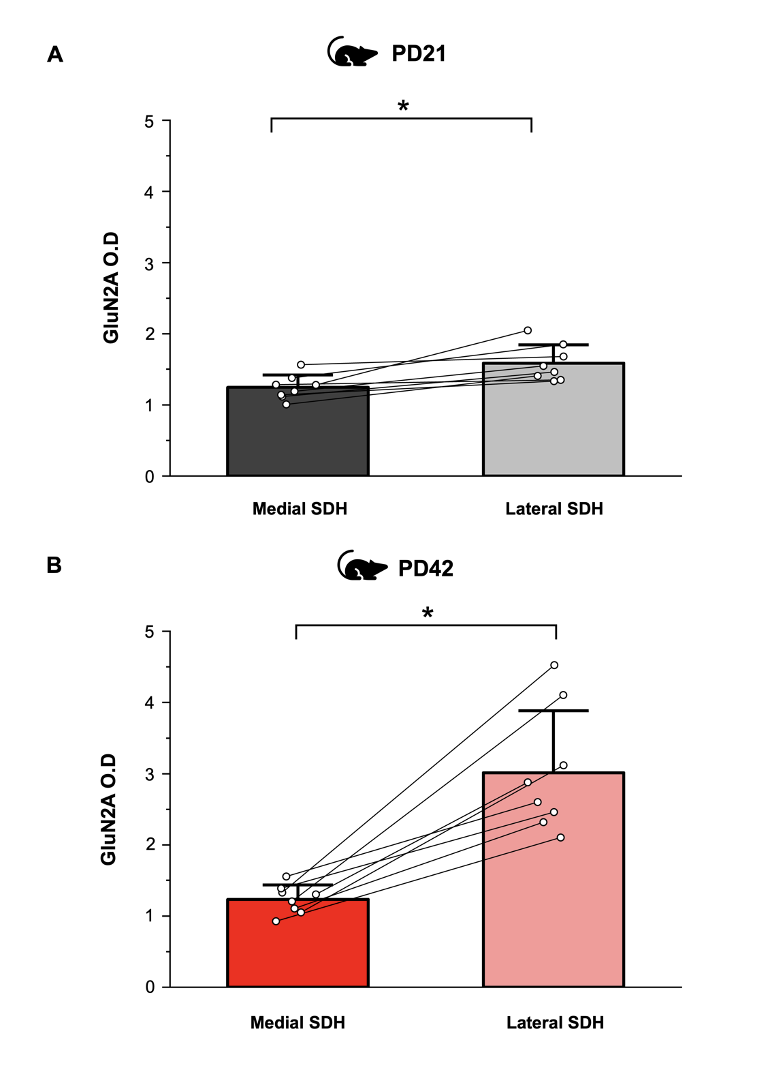
**Supplementary Figure 5. Asymmetrical distribution of GluN2A subunits with significantly heightened lateral SDH expression is minimal in juveniles and increases post puberty in adolescents for both sexes.** Quantitative analysis in **(A)** juvenile rats aged postnatal day 21 (*n* = 8 animals) and **(B)** adolescent rats aged postnatal day 42 (*n* = 8 animals) comparing the mean expression (normalized pixel intensity) for GluN2A subunits in the medial SDH versus the lateral SDH, with data combined across sexes. Data represents means ± SEM. *p<0.05.

**Supplementary Table 1. Statistic Summary Table**

| **Figure** | **Comparison** | **Statistical Test** | **Sample size (*n*)** | **Additional test statistics** | **p-value (2-tailed)** | **Significant?** |
| --- | --- | --- | --- | --- | --- | --- |
| 1A(v) | Adult Female Rat SDH vs DDH GluN2A | Paired samples t-test | 6 | SDH = 8.02 ± 1.37  DDH = 2.95 ± 0.45  *t*(5) = 5.25, Cohen's d = 2.15 | *p* = 0.0033 | Yes |
| 1B(v) | Adult Male Rat SDH vs DDH GluN2A | Paired samples t-test | 6 | SDH = 8.54 ± 1.10  DDH = 2.91 ± 0.25  *t*(5) = 5.44, Cohen's d = 2.22 | *p* = 0.0029 | Yes |
| 2A(v) | Adult Female Rat SDH vs DDH GluN2B | Paired samples t-test | 6 | SDH = 7.13 ± 1.16  DDH = 2.21± 0.23  *t*(5) = 4.88, Cohen's d = 1.99 | *p* = 0.0046 | Yes |
| 2B(v) | Adult Male Rat SDH vs DDH GluN2B | Paired samples t-test | 6 | SDH = 10.59 ± 1.72  DDH = 2.48 ± 0.31  *t*(5) = 5.54, Cohen's d = 2.26 | *p* = 0.0026 | Yes |
| 3A(v) | Adult Female Rat SDH vs DDH GluN2D | Paired samples t-test | 6 | SDH = 9.63 ± 2.66  DDH = 2.79 ± 0.26  *t*(5) = 2.71, Cohen's d = 1.12 | *p* = 0.042 | Yes |
| 3B(v) | Adult Male Rat SDH vs DDH GluN2D | Paired samples t-test | 6 | SDH = 5.99 ± 1.38  DDH = 2.32 ± 0.28  *t*(5) = 2.62, Cohen's d = 1.07 | *p* = 0.047 | Yes |
| 4A(v) | Adult Female Human SDH vs DDH GluN2A | Paired samples t-test | 12 | SDH = 5.40 ± 0.80  DDH = 2.39 ± 0.29  *t*(11) = 4.87, Cohen's d = 1.40 | *p* = 0.00050 | Yes |
| 4B(v) | Adult Male Human SDH vs DDH GluN2A | Paired samples t-test | 10 | SDH = 4.89 ± 0.84  DDH = 1.76 ± 0.27  *t*(9) = 4.30, Cohen's d = 1.36 | *p* = 0.0020 | Yes |
| 5A(v) | Adult Female Human SDH vs DDH GluN2B | Paired samples t-test | 12 | SDH = 4.80 ± 0.83  DDH = 2.00 ± 0.29  *t*(11) = 3.95, Cohen's d = 1.14 | *p*= 0.0023 | Yes |
| 5B(v) | Adult Male Human SDH vs DDH GluN2B | Paired samples t-test | 10 | SDH = 4.27 ± 1.15  DDH = 1.62 ± 0.23  *t*(9) = 2.67, Cohen's d = 0.84 | *p* = 0.026 | Yes |
| 6A(v) | Adult Female Human SDH vs DDH GluN2D | Paired samples t-test | 12 | SDH = 3.82 ± 0.38  DDH = 1.94 ± 0.24  *t*(11) = 5.89, Cohen's d = 1.70 | *p* = 0.00011 | Yes |
| 6B(v) | Adult Male Human SDH vs DDH GluN2D | Paired samples t-test | 10 | SDH = 4.26 ± 0.66  DDH = 1.73 ± 0.14  *t*(9) = 4.67, Cohen's d = 1.48 | *p* = 0.0012 | Yes |
| 7A(i) | Adult Rat SDH/DDH Ratio - Sex vs GluN2 Subunit | 2x3 Two-Way ANOVA | 6/group Total = 36 | Sex: *F*(1,30) = 0.401,  *partial η^2^* = 0.013 Subunit:  *F*(2,30) = 1.52, *partial η^2^* = 0.09 Sex * Subunit: *F*(2,30) = 1.16, *partial η^2^* = 0.07 | Sex: *p* = 0.53 Subunit: *p* = 0.24 Sex * Subunit:  *p* = 0.33 | Sex: No Subunit: No Sex * Subunit : No |
| 7A(ii) | Adult Human SDH/DDH Ratio - Sex vs GluN2 Subunit | 2x3 Two-Way ANOVA | Females: 12/group Males: 10/group Total observations = 66 | Sex: *F*(1,60) = 1.66,  *partial η^2^* = 0.03 Subunit:  *F*(2,60) = 0.812, *partial η^2^* = 0.026 Sex * Subunit: *F*(2,60) = 0.571, *partial η^2^* = 0.019 | Sex: *p* = 0.20 Subunit: *p* = 0.45 Sex * Subunit:  *p* = 0.57 | Sex: No Subunit: No Sex * Subunit : No |
| 7D(i) | Adult Rat Dorsal Horn qRT-PCR - Sex vs *Grin* Gene Target | 2x5 Two-Way ANOVA | Females: 8/group Males: 8/group Total observations = 80 | Sex: *F*(1,70) = 0.046,  *partial η^2^* = 6.53E-4 Gene Target:  *F*(4,70) = 191.26, *partial η^2^* = 0.92 Sex * Gene Target: *F*(4,70) = 0.39, *partial η^2^* = 0.02 | Sex: *p* = 0.83 Gene Target: *p* = 6.9E-37 Sex * Gene Target:  *p* = 0.82 | Sex: No Gene Target: Yes Sex * Gene Target: No |
| 7D(ii) | Adult Human Dorsal Horn qRT-PCR - Sex vs *Grin* Gene Target | 2x5 Two-Way ANOVA | Females: 8/group Males: 8/group Total observations = 80 | Sex: *F*(1,70) = 1.00,  *partial η^2^* = 0.01 Gene Target:  *F*(4,70) = 9.60, *partial η^2^* = 0.35 Sex * Gene Target: *F*(4,70) = 0.069, *partial η^2^* = 3.91E-3 | Sex: *p* = 0.32 Gene Target: *p* = 3.0E-6 Sex * Gene Target:  *p* = 0.99 | Sex: No Gene Target: Yes Sex * Gene Target: No |
| 8C(i) | Adult Rat Medial SDH/Lateral SDH Ratio - Sex vs GluN2 Subunit | 2x3 Two-Way ANOVA | Females: 6/group Males: 6/group Total observations = 36 | Sex: *F*(1,30) = 1.62,  *partial η^2^* = 0.05 Subunit:  *F*(2,30) = 2.45, *partial η^2^* = 0.14 Sex * Subunit: *F*(2,30) = 0.516, *partial η^2^* = 0.033 | Sex: *p* = 0.21 Subunit: *p* = 0.10 Sex * Subunit:  *p* = 0.60 | Sex: No Subunit: No Sex * Subunit : No |
| 8C(ii) | Adult Human Medial SDH/Lateral SDH Ratio - Sex vs GluN2 Subunit | 2x3 Two-Way ANOVA | Females: 12/group Males: 10/group Total observations = 66 | Sex: *F*(1,60) = 0.662,  *partial η^2^* = 0.011 Subunit:  *F*(2,60) = 1.94, *partial η^2^* = 0.06 Sex * Subunit: *F*(2,60) = 0.635, *partial η^2^* = 0.021 | Sex: *p* = 0.42 Subunit: *p* = 0.15 Sex * Subunit:  *p* = 0.53 | Sex: No Subunit: No Sex * Subunit : No |
| 10C | Rat SDH/DDH GluN2A Ratio - Sex vs Age | 2x3 Two-Way ANOVA | PD21: 4/group PD42: 4/group PD90+: 6/group Total observations = 28 | Sex: *F*(1,22) = 0.017,  *partial η^2^* = 0.001 Age:  *F*(2,22) = 19.4,  *partial η^2^* = 0.6 Sex * Age: *F*(2,22) = 0.971, *partial η^2^* = 0.081 | Sex: *p* = 0.90 Age: *p* = 1.4E-5 Sex * Age:  *p* = 0.39 | Sex: No Age: Yes Sex * Age : No |
| 10D | Rat Medial SDH/Lateral SDH GluN2A Ratio - Sex vs Age | 2x3 Two-Way ANOVA | PD21: 4/group PD42: 4/group PD90+: 6/group Total observations = 28 | Sex: *F*(1,22) = 0.002,  *partial η^2^* = 0.000 Age:  *F*(2,22) = 9.88,  *partial η^2^* = 0.47 Sex * Age: *F*(2,22) = 1.756, *partial η^2^* = 0.138 | Sex: *p* = 0.97 Age: *p* = 0.00087 Sex * Age:  *p* = 0.20 | Sex: No Age: Yes Sex * Age : No |
| 10E | Juvenile vs Adult mEPSC Charge Transfer in Lamina II Neurons | Independent samples t-test | PD20-22: 5 neurons PD90+: 11 neurons | PD20-22 Charge Transfer = 1.31 ± 0.39 pC PD90+ Charge Transfer =  2.50 ± 0.22 pC | *p* = 0.011 | Yes |
| Suppl. 1A | Adult Female Rat Medial SDH vs Lateral SDH GluN2A | Paired samples t-test | 6 | Medial SDH = 1.89 ± 0.28  Lateral SDH = 4.36 ± 0.85  *t*(5) = -3.56, Cohen's d = -1.45 | *p* = 0.016 | Yes |
| Suppl. 1B | Adult Female Rat Medial SDH vs Lateral SDH CGRP | Paired samples t-test | 6 | Medial SDH = 1.07 ± 0.06  Lateral SDH = 1.01 ± 0.11  *t*(5) = 0.75, Cohen's d = 0.31 | *p* = 0.49 | No |
| Suppl. 2A | Adult Rat Thoracic Medial SDH vs Lateral SDH GluN2A (Sexes Combined) | Paired samples t-test | 6 | Medial SDH = 1.48 ± 0.19  Lateral SDH = 2.07 ± 0.39  *t*(5) = -2.77, Cohen's d = -1.13 | *p* = 0.039 | Yes |
| Suppl. 2B | Adult Rat Thoracic Medial SDH vs Lateral SDH NeuN (Sexes Combined) | Paired samples t-test | 6 | Medial SDH = 1.03 ± 0.07  Lateral SDH = 0.99 ± 0.12  *t*(5) = 0.43, Cohen's d = 0.18 | *p* = 0.68 | No |
| Suppl. 3A(i) | PD21 Female Rat SDH vs DDH GluN2A | Paired samples t-test | *4* | SDH = 2.75 ± 0.70  DDH = 1.96 ± 0.31 *t*(3) = 1.67, Cohen's d = 0.83 | *p* = 0.19 | No |
| Suppl. 3A(ii) | PD21 Male Rat SDH vs DDH GluN2A | Paired samples t-test | 4 | SDH = 2.90 ± 0.62  DDH = 1.92 ± 0.39  *t*(3) = 3.13, Cohen's d = 1.57 | *p* = 0.052 | No |
| Suppl. 3B(i) | PD42 Female Rat SDH vs DDH GluN2A | Paired samples t-test | 4 | SDH = 8.35 ± 1.61  DDH = 2.66 ± 0.50  *t*(3) = 4.91, Cohen's d = 2.46 | *p* = 0.016 | Yes |
| Suppl. 3B(ii) | PD42 Male Rat SDH vs DDH GluN2A | Paired samples t-test | 4 | SDH = 6.38 ± 0.72  DDH = 2.27 ± 0.16  *t*(3) = 6.64, Cohen's d = 3.32 | *p* = 0.0070 | Yes |
| Suppl. 4A | PD21 Rat Medial SDH vs Lateral SDH GluN2A (Sexes Combined) | Paired samples t-test | 8 | Medial SDH = 1.25 ± 0.06  Lateral SDH = 1.59 ± 0.09  *t*(7) = -4.31, Cohen's d = -1.52 | *p* = 0.0035 | Yes |
| Suppl. 4B | PD42 Rat Medial SDH vs Lateral SDH GluN2A (Sexes Combined) | Paired samples t-test | 8 | Medial SDH = 1.24 ± 0.07 = Lateral SDH = 3.02 ± 0.31 = *t*(7) = -5.90, Cohen's d = -2.08 | *p* = 0.00060 | Yes |
